# Lasting aversive consequences of a single dose of 4-hydroxytamoxifen

**DOI:** 10.64898/2026.09.16.751757

**Authors:** Evan J. Kyzar, Ruby Setara, Maya Eisengart, Sophia Virkar, Lydia Rogerson, Alejandro Ramirez, C. Daniel Salzman

**Affiliations:** Department of Neuroscience, Columbia University, New York, NY, USA; The Mortimer B. Zuckerman Mind Brain Behavior Institute, Columbia University, New York, NY, USA; Department of Psychiatry, Columbia University, New York, NY, USA; New York State Psychiatric Institute, New York, NY, USA; Kavli Institute for Brain Science, Columbia University, New York, NY, USA

## Abstract

4-hydroxytamoxifen (4-OHT) is commonly used to facilitate activity-dependent recombination in neurons. Here we show that one dose of 4-OHT devalues alcohol, contexts, and tastants, indicating that experimental designs using 4-OHT should control for its aversiveness. Further, reactivation of 4-OHT-responsive neurons causes conditioned taste avoidance. An activity screen coupled with graph theoretical analysis identifies candidate brain areas mediating these effects. We suggest these areas could be therapeutic targets for treating addiction.

---

The development of tools for tagging and perturbing the activity of neural ensembles based on their activity has greatly advanced neuroscientific investigations into the neural basis of learning and memory, motivational states, and addiction^1–10^. Typically, genetically modified mice (e.g., FosTRAP and Trap2 mice^3,6,10^) are employed whereby high doses of estrogen receptor modulators, such as tamoxifen or its active metabolite 4-hydroxytamoxifen (4-OHT)^1–7^ are required to drive activity-dependent recombination. Estrogen receptor modulators are known to exert behavioral effects independent of their use in driving activity-dependent recombination^11–14^. In this paper, we test how 4-OHT delivery impacts the behavioral assessment of different types of stimuli and situations, including during alcohol and taste preference assays and during place avoidance.

To study the development and expression of alcohol preference, we initially performed a two-bottle choice (2BC) alcohol drinking task in Trap2:Ai14 mice (**Extended Data Fig. 1A**). Strikingly, a single dose of 4-OHT led to a marked decrease in alcohol preference and consumption which persisted for multiple weeks (**Extended Data Fig. 1B,C**); total fluid intake was altered only for the first two weeks (**Extended Data Fig. 1D**).

To determine if this effect was specific to Trap2 mice, we performed an identical assay in C57BL/6J mice but with a vehicle control (**Fig. 1A**). The same dose of 4-OHT used to induce recombination in Trap2 mice (50mg/kg) led to decreased alcohol preference and consumption in C57BL/6J mice compared to vehicle-exposed controls (**Fig. 1B,C**). This effect persisted for weeks without substantive changes in total fluid consumption or body weight (**Extended Data Fig. 2A,B**). Females resumed higher alcohol preference and consumption more rapidly following 4-OHT exposure, consistent with the fact that female mice typically consume more alcohol relative to males^15,16^ (**Extended Data Fig. 2C-F**).

**Figure 1.**
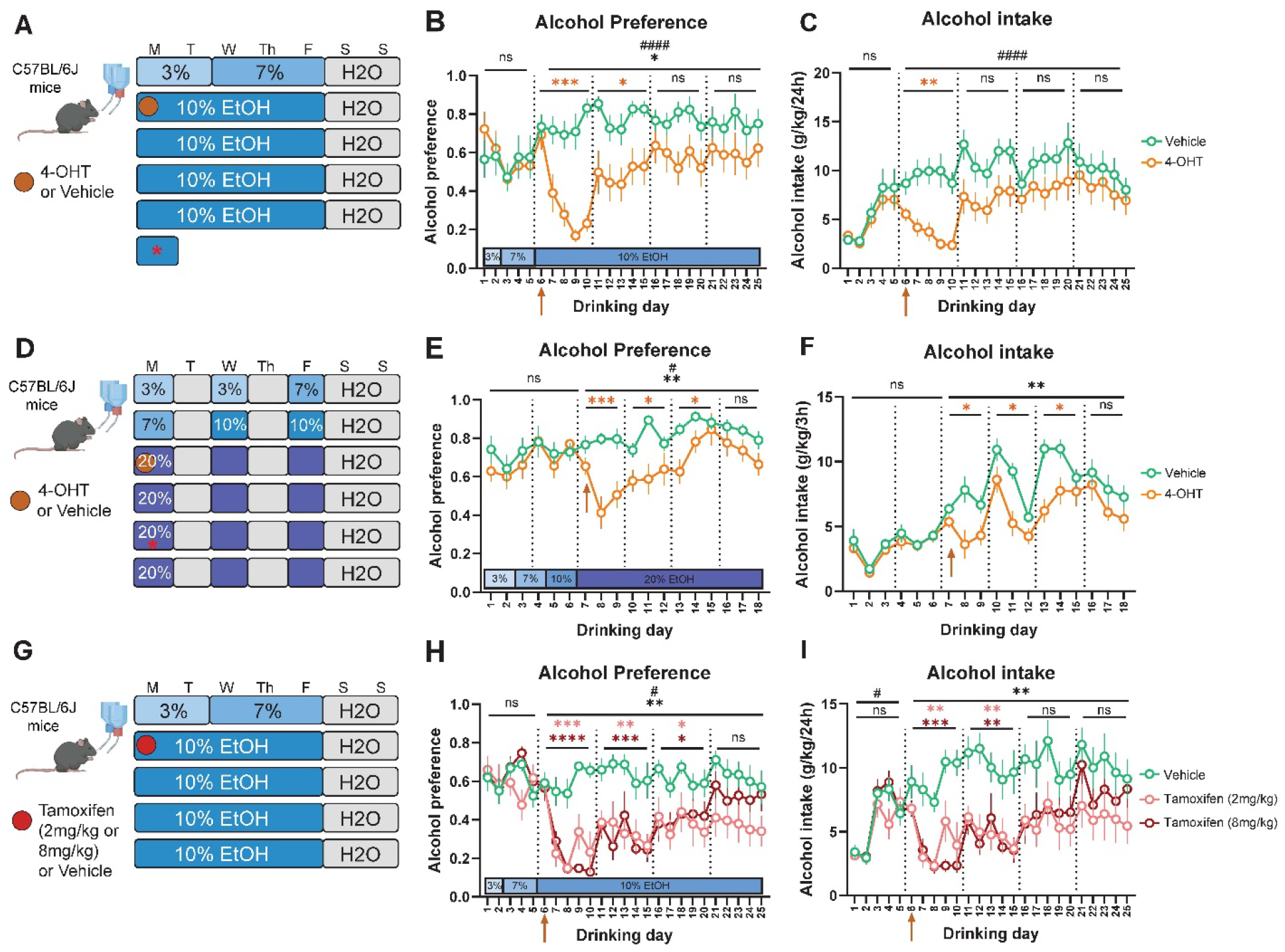
A single dose of 4-hydroxytamoxifen (4-OHT) or tamoxifen leads to decreased alcohol preference. A) Experimental schematic of exposure of C57BL/6J mice to 4-OHT (50mg/kg; i.p.) or vehicle on the first day of 10% alcohol exposure. Mice were given continuous access to alcohol for 5d followed by a 2d washout period each week. Red asterisk indicates day of blood collection. See Methods for blood alcohol concentrations. B) Alcohol preference (calculated as mL of alcohol consumed per day divided by total fluid consumed per day) in C57BL/6J mice exposed to 4-OHT compared to vehicle as shown in A). 2-way RM-ANOVA omnibus effects: *F_Group_(1,14) = 8.793, *p*=0.01; ^####^F_GroupxTime_(19,266) = 3.804, *p*<0.0001. No significant differences between groups in pre-treatment period. Arrow indicates day of 4-OHT/Vehicle treatment, and dotted lines separate weekly averages of preference used to calculate effect persistence. Orange asterisks represent differences between groups in the weekly rolling average of alcohol preference by multiplicity-corrected Šidák tests after 2-way RM ANOVA. \*\*\**p*<0.001; \**p*<0.05. n=8. C) Alcohol intake (g/kg/24h) in C57BL/6J mice exposed to 4-OHT compared to vehicle as shown in A). 2-way RM-ANOVA omnibus effects: F_Group_(1,14) = 3.754, *p*=0.07; ^####^F_GroupxTime_(19,266) = 3.047, *p*<0.0001. No significant differences between groups in pre-treatment period. Arrow indicates day of 4-OHT/Vehicle treatment, and dotted lines separate weekly averages of preference used to calculate effect persistence. Orange asterisks represent differences between groups in the weekly rolling average of alcohol intake by multiplicity-corrected Šidák tests after 2-way RM ANOVA. \*\**p*<0.01. n=8. D) Experimental schematic of exposure of C57BL/6J mice to 4-OHT (50mg/kg; i.p.) or vehicle on the first day of 20% alcohol exposure in an intermittent access 2BC where mice were given access to alcohol for 3hrs on Mondays, Wednesdays, and Fridays each week. Red asterisk indicates day of blood collection. See Methods for blood alcohol concentrations. E) Alcohol preference in C57BL/6J mice exposed to 4-OHT compared to vehicle as shown in D). 2-way RM-ANOVA omnibus effects: **F_Group_(1,14) = 15.90, *p*=0.0013; ^#^F_GroupxTime_(5.940, 83.16) = 2.360, *p*=0.038. No significant differences between groups in pre-treatment period. Arrow indicates day of 4-OHT/Vehicle treatment, and dotted lines separate weekly averages of preference used to calculate effect persistence. Orange asterisks represent differences between groups in the weekly rolling average of alcohol preference by multiplicity-corrected Šidák tests after 2-way RM ANOVA. \*\*\**p*<0.001; \**p*<0.05. n=8. F) Alcohol intake (g/kg/24h) in C57BL/6J mice exposed to 4-OHT compared to vehicle as shown in D). 2-way RM-ANOVA omnibus effects: **F_Group_(1,14) = 14.05, *p*=0.0022; F_GroupxTime_(6.163, 86.28) = 1.540, *p*=0.17. No significant differences between groups in pre-treatment period. Arrow indicates day of 4-OHT/Vehicle treatment, and dotted lines separate weekly averages of preference used to calculate effect persistence. Orange asterisks represent differences between groups in the weekly rolling average of alcohol intake by multiplicity-corrected Šidák tests after 2-way RM ANOVA. \**p*<0.05. n=8. G) Experimental schematic of exposure of C57BL/6J mice to tamoxifen (2mg/kg or 8mg/kg; i.p.) or vehicle on the first day of 10% alcohol exposure. Mice were given continuous access to alcohol for 5d followed by a 2d washout period each week. H) Alcohol preference in C57BL/6J mice exposed to tamoxifen compared to vehicle as shown in G). 2-way RM-ANOVA omnibus effects: *\*\**F_Group_(2,20) = 9.864, *p*=0.001; ^#^F_GroupxTime_(13.52, 135.2) = 1.851, *p*=0.039. No significant differences between groups in pre-treatment period. Arrow indicates day of tamoxifen/Vehicle treatment, and dotted lines separate weekly averages of preference used to calculate effect persistence. Asterisks represent differences between groups (light red: significant in vehicle vs 2mg/kg tamoxifen; dark red: significant in vehicle vs 8mg/kg tamoxifen) in the weekly rolling average of alcohol preference by multiplicity-corrected Šidák tests after 2-way RM ANOVA. \*\*\*\**p*<0.0001; \*\*\**p*<0.001; \*\**p*<0.01; \**p*<0.05. n=7-8. I) Alcohol intake (g/kg/24h) in C57BL/6J mice exposed to tamoxifen compared to vehicle as shown in G). 2-way RM-ANOVA omnibus effects: *\*\**F_Group_(2,20) = 6.988, *p*=0.005; F_GroupxTime_(10.93, 109.3) = 1.445, *p*=0.16. No significant differences between groups in pre-treatment period (group x time interaction omnibus reached significance but there were no differences in daily alcohol intake between any groups). Arrow indicates day of tamoxifen/Vehicle treatment, and dotted lines separate weekly averages of preference used to calculate effect persistence. Asterisks represent differences between groups (light red: significant in vehicle vs 2mg/kg tamoxifen; dark red: significant in vehicle vs 8mg/kg tamoxifen) in the weekly rolling average of alcohol intake by multiplicity-corrected Šidák tests after 2-way RM ANOVA. \*\*\**p*<0.001; \*\**p*<0.01. n=7-8.

4-OHT delivery also diminished alcohol preference in other alcohol access schedules. In a modified intermittent binge-access (IA2BC) paradigm (**Fig. 1D**), 4-OHT exposure on the first day of 20% alcohol access led to a three-week decrease in preference and intake (**Fig. 1E,F; Extended Data Fig. 3A,B**) without affecting total fluid intake (**Extended Data Fig. 3C,D**). In a continuous access 2BC test (**Extended Data Fig. 3E**), mice exposed to 4-OHT demonstrated altered alcohol preference and intake and consumption (**Extended Data Fig. 3F-K**).

Notably, 4-OHT did not diminish alcohol preference in all situations. Exposure to 4-OHT after 4 weeks of consuming alcohol did not substantively alter preference or consumption (**Extended Data Fig. 3L-O**), suggesting that alcohol preference is more difficult to diminish after its preference is well-established. If 4-OHT was not delivered in association with drinking alcohol, such as prior to alcohol initiation in a 2BC task, alcohol preference and consumption also were not altered (**Extended Data Fig. 4A-D**).

The effect of 4-OHT was also observed using doses as low as 1mg/kg delivered to female mice, indicating that behavioral effects may emerge well below doses used to drive recombination (**Extended Data Fig. 4E,F**). Further, a single dose of tamoxifen at two clinically relevant doses (2mg/kg & 8mg/kg) decreased alcohol preference and intake for weeks without altering total fluid intake (**Fig. 1G-I; Extended Data Fig. 4G-J**). Of note, the half-life of tamoxifen is 12hrs compared to 4-6hrs for 4-OHT^14,17^. Tamoxifen is processed through conversion to 4-OHT, and this prolonged exposure might explain why decreased alcohol preference persists longer in tamoxifen-treated mice than in 4-OHT-treated mice.

We hypothesized that the diminished preference for alcohol arose because 4-OHT was acting as if it was an aversive unconditioned stimulus. Under this hypothesis, pairing 4-OHT delivery with alcohol consumption leads to devaluation and subsequent avoidance. If this hypothesis is correct, then 4-OHT delivery should have analogous effects on other assays. The effects of 4-OHT delivery on conditioned taste avoidance (CTA), conditioned place avoidance (CPA), and a sucrose preference task (SPT) were therefore tested.

To probe conditioned taste avoidance, we habituated animals to a 2-bottle setup for 2 hours daily prior to pairing 4-OHT (50mg/kg; given 30 min after end of session) or vehicle with a flavored tastant (**Fig. 2A**). Typically, CTA is only observed if the conditioned flavor is novel (i.e., mice have not encountered it prior to the conditioning session). For example, lithium chloride (LiCl), a compound commonly used for aversive conditioning, induces CTA for novel but not familiar flavors^18^. In our experiments, when a flavored tastant was novel on the conditioning day, mice subsequently avoided that flavor following 4-OHT pairing (**Fig. 2B**). Notably, 4-OHT also caused avoidance when paired with a familiar flavor that mice were exposed to for 2 days prior to pairing (**Fig. 2C**). In a separate CTA experiment, we compared 4-OHT delivery to LiCl with respect to avoidance of a novel flavor, observing effects of similar magnitude (**Extended Data Fig. 5A**). A single dose of LiCl also altered alcohol preference in a 2BC task but did not decrease weekly average preference or intake (**Extended Data Fig. 5B-H**).

**Figure 2.**
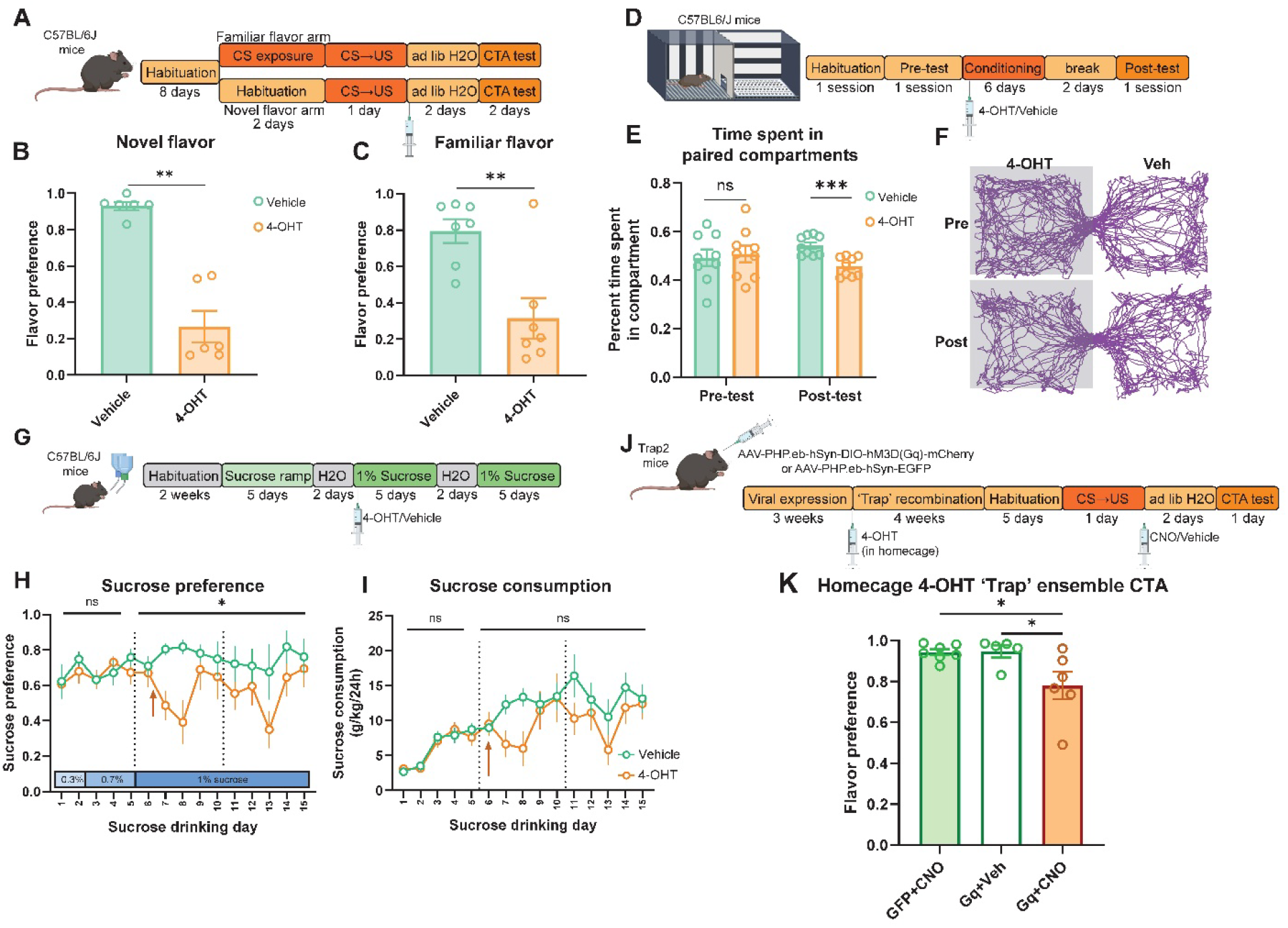
4-OHT causes avoidance of paired contexts and tastants. A) Conditioned taste avoidance (CTA) experimental schematic. Mice were habituated to drinking water from a two-bottle setup while fasted in 2hr sessions (only water available) for 8 days. Mice given 4-OHT or vehicle in conjunction with a novel flavor (0.3% saccharin in 0.06% Kool-Aid powder, also known as the conditioned stimulus or CS) were given an additional 2 water habituation sessions, while mice exposed to Kool-Aid as a familiar flavor were given 2 sessions of Kool-Aid consumption. All habituation sessions were followed by sham saline injections. On the conditioning day, mice received either 4-OHT (50mg/kg i.p.; as the unconditioned stimulus or US) or vehicle 30 min after the 2hr consumption session. Mice were given 2d of *ad libitum* water access prior to a 2BC test to assess flavor preference. B) Flavor preference for mice exposed to 4-OHT or Vehicle in conjunction with a novel flavor. Mann-Whitney *U*=0, *p*=0.002. n=6. C) Flavor preference for mice exposed to 4-OHT or Vehicle in conjunction with a familiar flavor. Unpaired *t*-test; *t*(12)=3.709, *p*=0.003. n=7. D) Conditioned place avoidance (CPA) experimental schematic. Mice were habituated to a two-compartment place preference apparatus prior to a pre-test. Mice received 6 unbiased conditioning days (3 each for 4-OHT or vehicle) where they explored one compartment for 2hr after a 4-OHT (50-mg/kg; i.p.) or vehicle injection. Place avoidance/preference was then tested in a 10min ‘post-test’ free exploration session. E) Time spent in paired compartments in pre-test vs post-test sessions. Pre: *t*(16)= 0.3171, *p*=0.76. Post: *t*(16)= 4.47, *p*=0.0004. Unpaired *t*-tests. n=9. F) Representative trace of pre- and post-test exploration pattern in a single mouse. G) Experimental schematic of sucrose preference test (SPT) where C57BL/6J mice were exposed to 4-OHT (50mg/kg; i.p.) or vehicle on the first day of 1% sucrose exposure. Mice were given continuous access to sucrose for 5d followed by a 2d washout period each week. H) Sucrose preference in C57BL/6J mice exposed to 4-OHT compared to vehicle as shown in G). 2-way RM-ANOVA omnibus effects: *F_Group_(1,15) = 4.630, *p*=0.048; F_GroupxTime_(9, 135) = 1.228, *p*=0.28. No significant differences between groups in pre-treatment period. Arrow indicates day of 4-OHT/Vehicle treatment, and dotted lines separate weekly averages of preference used to calculate effect persistence. No significant differences between groups in the weekly rolling average of sucrose preference by multiplicity-corrected Šidák tests after 2-way RM ANOVA (see Methods for note on conservative *post-hoc* testing). n=8-9. I) Sucrose consumption (g/kg/24h) in C57BL/6J mice exposed to 4-OHT compared to vehicle as shown in G). 2-way RM-ANOVA omnibus effects: F_Group_(1,15) = 1.433, *p*=0.25; F_GroupxTime_(9, 135) = 1.741, *p*=0.09. No significant differences between groups in pretreatment period. Arrow indicates day of 4-OHT/Vehicle treatment, and dotted lines separate weekly averages of preference used to calculate effect persistence. No significant differences between groups in the weekly rolling average of sucrose consumption by multiplicity-corrected Šidák tests after 2-way RM ANOVA. n=8-9. J) Experimental schematic for assessing the aversiveness of a homecage 4-OHT ‘Trap’ ensemble. Trap2 mice were given a retroorbital injection of a CNS-penetrant chemogenetic construct (AAV-PHP.eb-hSyn-DIO-hM3D(Gq)-mCherry^19^; Gq+Veh & Gq+CNO groups) or control virus (AAV-PHP.eb-hSyn-EGFP; GFP+CNO group). Mice were allowed to recover for 3 weeks prior to activation of a homecage ‘Trap’ ensemble via injection of 4-OHT (50mg/kg; i.p.). After 4 weeks, mice were habituated to drinking water from a two-bottle setup while fasted in 2hr sessions (only water available) for 5 days. All habituation sessions were followed by sham saline injections. On the conditioning day, mice received either CNO (5mg/kg) or vehicle 30 min after a 2hr Kool-Aid consumption session. Mice were given 2d of *ad libitum* water access prior to a 2BC test to assess flavor preference. K) Flavor preference for mice exposed to CNO or Vehicle in conjunction with a novel flavor as shown in J). One-way ANOVA with Tukey *post-hoc* tests between groups: F_Group_(2, 15) = 5.032, *p*=0.021; *p*=0.99 (GFP+CNO vs. Gq+Veh), *p*=0.033 (GFP+CNO vs. Gq+CNO), *p*=0.044 (Gq+Veh vs. Gq+CNO). n=5-7.

To determine if 4-OHT induces conditioned place avoidance (CPA), we paired a mouse’s presence in particular compartments of a place preference/avoidance box with either vehicle or 4-OHT delivery (**Fig. 2D**). Mice did not have a compartment preference prior to conditioning but spent significantly less time in the 4-OHT-paired compartment after conditioning (**Fig. 2E,F**; **Extended Data Fig. 6A-C**). In a sucrose preference task, mice were given a choice of two bottles in the homecage – one filled with water and the other with increasing concentrations of sucrose (**Fig. 2G**). Exposure to 4-OHT on the first day of 1% sucrose exposure altered sucrose preference though *post-hoc* weekly averages did not meet multiplicity-corrected significance criteria (**Fig. 2H,I; Extended Data Fig. 7A,B**).

We considered the possibility that a nonspecific effect of 4-OHT on locomotion could lead to altered preference behavior. We therefore assayed mouse behavior in an unconstrained open field test and found no effect of 4-OHT on center time, distance traveled, or common behaviors including grooming and rearing compared to vehicle-treated mice (50mg/kg and 1mg/kg doses 30 min prior to testing; **Extended Data Fig. 7C-D**).

In typical experiments using activity dependent expression to test the functional role of neurons based on their physiological response properties, 4-OHT is administered and then mice undergo a particular experience, such as the presentation of a stimulus. The assumption is that the neurons activated by the stimulus are those that will be trapped, and that subsequent perturbation experiments (e.g. with opto- or chemogenetic approaches) can test the causal role of those neurons on a specific behavior. However, the aversive effects of 4-OHT across multiple assays raises the possibility that the actions of 4-OHT itself may activate, and therefore ‘trap’, neurons. Recombination in these ‘trapped’ neurons would be unrelated to the intended behavioral assay. If this occurs, behavioral results may result from perturbations of neurons activated by 4-OHT and not to the intended experimental parameter. We therefore designed an experiment to determine if perturbing the activity of neurons responsive to 4-OHT delivery affects behavior.

Trap2 mice were given a peripheral injection of a virus carrying a chemogenetic construct that penetrates the CNS (AAV-PHP.eb-hSyn-DIO-hM3D(Gq)-mCherry; or control eGFP vector)^19^. 4-OHT was delivered while mice were in their homecage in the absence of any specific stimulus. Subsequently, we trained mice on a CTA. Different groups of mice were given either CNO or vehicle following exposure to a novel flavor (**Fig. 2J**). Activating the ‘trapped’ ensemble was sufficient to decrease flavor preference compared to vehicle-exposed Trap2 mice as well as CNO-exposed control vector mice (**Fig. 2K**).

The ability of a single dose of 4-OHT to cause a rapid and sustained decrease in alcohol preference raises the possibility of using this capability to understand the neural mechanisms of stimulus devaluation. Most assays that induce alcohol devaluation rely on changing the properties of alcohol as a stimulus (e.g., adulterating alcohol with quinine^20,21^). 4-OHT, on the other hand, may be acting by activating particular neurons that drive aversively motivated behavior, consistent with the experiments showing that re-activating neurons responsive to 4-OHT diminishes flavor preference.

As a first step towards identifying brain areas that mediate how 4-OHT diminishes alcohol preference, we utilized a recently developed neural activity screening pipeline, two-timepoint statistical inference with subtraction (TTP-S)^5^. Trap2:Ai14 mice received 4-OHT on the first day of 10% alcohol drinking and were sacrificed 1 week later for c-Fos staining after re-exposure to alcohol (EE group). Another group of mice received 4-OHT at the same timepoint, but mice were sacrificed for c-Fos staining while only drinking water (EH group) (**Fig. 3A**). Cells double-labeled in the EH group are presumed to lack specificity to alcohol exposure since those cells would have been active both during alcohol exposure and in the during water-only access (**Fig. 3B**).

**Figure 3.**
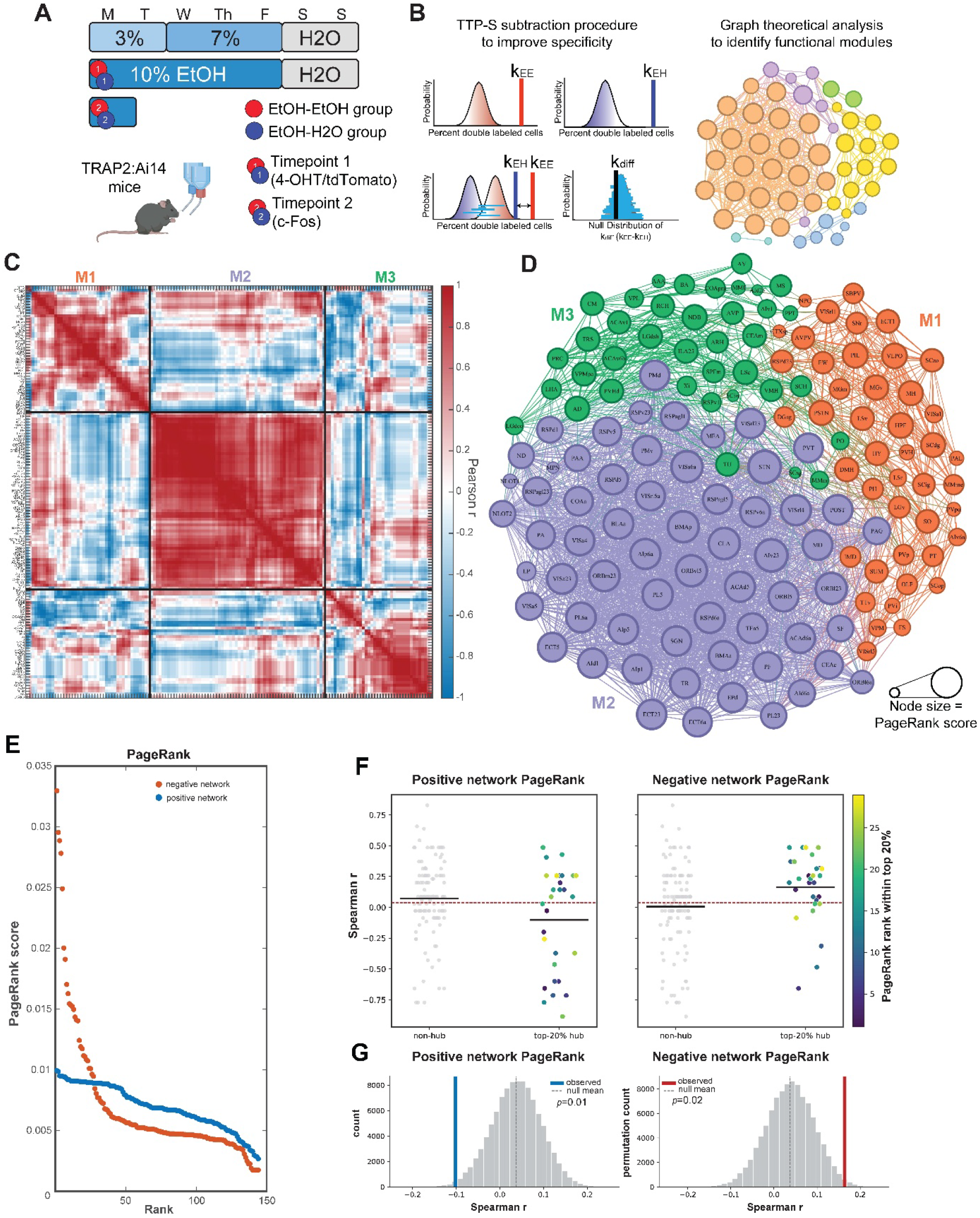
Whole-brain activation paflerns in 4-OHT-mediated devaluation. A) Experimental schematic of two-timepoint design comparing Trap2xAi14 mice drinking alcohol in a two-bottle choice (2BC) task. Mice in ethanol-ethanol (EE) group were given 4-OHT (50mg/kg; i.p.) on the first day of 10% alcohol exposure (to devalue alcohol) followed by sacrifice for c-Fos staining 1 week later upon re-exposure to alcohol. Another group of mice (ethanol-H2O; EH group) was given 4-OHT (50mg/kg) on the first day of 10% alcohol exposure followed by sacrifice for c-Fos staining 1 week later while only having access to water. The EH group was used for the subtraction procedure (see Methods) B) Two-timepoint statistical inference with subtraction (TTP-S) workflow^5^. Immunohistochemistry and whole-brain imaging were performed on mice from A). Images were registered to the Allen Brain Atlas Common Coordinate Framework, and automated cell counting generated *p*-values for each brain area indicating the potential over-representation of double-labeled cells (at both timepoint 1; tdTomato & timepoint 2; c-Fos). The double-labeled cell counts from the EH group were “subtracted” (see Methods) from the EE counts to correct for neurons demonstrating high baseline activity that may represent false positives. Graph theoretical analysis was then performed to identify hub areas. See Methods for further details. C) Correlation matrix of counts of neurons active at both timepoints in the alcohol devaluation (EE) group (n=5). Hierarchical clustering determined the location of brain areas on the matrix, and modules were determined using Louvain averaging over 500 iterations. Note that brain regions are represented in the same order in columns and rows. D) Positively-correlated network connectivity in the alcohol devaluation (EE) group (n=5). Each line indicates a positive correlation (R>0.6) between brain areas [Fruchterman-Reingold layout]; Colored nodes indicate module assignment, and the size of the node indicates the PageRank score. E) PageRank score to identify hub brain regions in negative (red) and positive (blue) correlation matrices in alcohol devaluation (EE) group. F) Correlation values from hub brain regions from alcohol devaluation (EE) applied to data from Fig. 2J-L. Left) The top 20% of regions by PageRank in the positive correlation matrix show a more negative correlation with flavor preference (Spearman *r* = -0.102) than expected by chance. The other 80% of regions are shown in grey for comparison. Right) The top 20% of regions by PageRank in the negative correlation matrix show a more negative correlation with flavor preference (Spearman *r* = 0.164) than expected by chance. The other 80% of regions are shown in grey for comparison. Colors indicate rank from 1-29 for top 20% of regions by PageRank. G) Estimation plots for expected and null distribution for data in F). Regions were sampled 100,000 times to create a null distribution, with observed data shown in blue (left; positive) and red (right; negative). *p*-values indicate comparison to null distribution by permutation test corrected for multiple comparisons.

One hundred and seventy-five brain areas met multiple comparisons-corrected significance for overlap in activation between the two timepoints in the EE group (**Table S1).** Graph theoretical analysis revealed three tightly correlated modules of double-labeled brain regions (**Fig. 3C,D; Extended Data Fig. 8A**). Additionally, the modular correlation structure of EE ‘alcohol devaluation’ mice was distinct from EH mice, from mice exposed to both timepoints in the homecage, and from mice exposed to 4-OHT after 6 weeks of alcohol consumption (**Extended Data Fig. 8B-D**). Enrichment analysis demonstrated that Module 1 was enriched for hypothalamic regions; Module 2 was enriched for cortical and cortical subplate regions (i.e., basolateral [BLA] and basomedial amygdala [BMA]); and Module 3 was enriched for thalamic nuclei (**Extended Data Fig. 8E,F; Table S2**). Notably, Module 2 included many areas implicated in aversive learning and stimulus valuation including BLA/BMA, insular, orbitofrontal, and anterior cingulate areas^22–25^. Module 3 contained the classic Papez circuit consisting of the mammillary bodies and anterior thalamus implicated in episodic memory^26,27^.

We used the PageRank measure from graph theoretical analysis to identify putative hub regions^5,28^ in both the positive and negative correlation networks (**Fig. 3E**). We reasoned that regions with high PageRank scores would be more likely to be correlated with avoidance-related behavior, accounting for 4-OHT-induced devaluation. To test this, we explored the correlation of PageRank values from the devaluation TTP-S analysis with the correlation between 4-OHT-activated cells and flavor preference from the homecage ‘Trap’ CTA experiment in **Fig. 2J,K**. We trained a custom model^29,30^ to measure 4-OHT-activated cells across the brain. Note that there was no correlation between total activated cells and flavor preference. Moreover, no brain areas met FDR-corrected significance for correlation with preference (**Table S3**, **Extended Data Fig. 9A-G**; 12 individual brain regions were significantly correlated with preference at nominal *p*<0.05 levels that did not survive FDR-corrected significance).

The critical question is whether the activity screen coupled with graph theoretical analysis identifies brain areas that could play a role in devaluation. For each brain area, we first computed the Spearman *r* value that quantifies the correlation across mice between the number of 4-OHT ‘trapped’ cells with flavor preference. Next, we asked whether the PageRank score for a brain area identified as activated in the TTP-S screen predicts its Spearman *r* value. We discovered that positive network PageRank scores were negatively correlated with the Spearman *r* values quantifying the relationship between the number of 4-OHT ‘trapped’ cells with flavor preference; by contrast, negative network PageRank scores were positively correlated with the same Spearman *r* values (**Extended Data Fig. 9H,I; Table S4**). This effect was most prominent when examining the 20% of brain regions (hubs) with the highest PageRank scores (**Fig. 3F**). Hubs identified in the positive network had lower correlation values than expected by chance, whereas hubs from the negative network had significantly higher correlation values (positive network; **Fig. 3G, left**; negative network, **Fig. 3G, right**). Thus, the activation of hub regions from the positive network (regions positively correlated with many other significant regions) is associated with greater flavor avoidance in a CTA, whereas activation of negative hub regions is associated with less avoidance. These data reveal that by combining TTP-S with graph theoretical analysis, brain areas likely to play a mechanistic role in devaluation can be identified. Approaches that only examine the number of cells activated by 4-OHT in each brain area would fail to identify these brain areas with this sample size.

Our results indicate that 4-OHT and tamoxifen delivery have aversive consequences that can endure for weeks. When delivery is paired with otherwise rewarding substances, such as alcohol and sucrose, preference for these substances is diminished. When paired with a particular place, mice avoid that place. Finally, when neurons activated by 4-OHT are re-activated chemogenetically, preference for a desirable tastant is diminished.

The strong aversive effects of 4-OHT have critical implications for experimental design and interpretation given the use of the Trap2 mouse model and similar methods for activity-dependent recombination^1–7^. Re-activating an ensemble ‘trapped’ by 4-OHT delivery may cause effects on behavior due to neurons activated only by 4-OHT, and/or due to changes in neural activity related to the aversive consequences of 4-OHT delivery when paired with an experimentally selected stimulus, action, or experience. Consistent with our findings, prior studies have observed that re-activating neural ensembles tagged by 4-OHT tends to lead to aversive or anxiogenic behavioral effects much more frequently than appetitive or anxiolytic effects (**Extended Data Fig. 10).** We suggest that experimental designs using 4-OHT to induce recombination should therefore include additional control groups, such as groups of mice where 4-OHT is delivered either in the homecage, or in association with additional “control” stimuli or events. The TTP-S pipeline is one example, as stimulus exposure occurs at two timepoints, one of which does not require 4-OHT treatment, and the overlap between neural activity is examined^5^. Alternatively, one could adopt methods of neuronal tagging that do not require 4-OHT delivery, such as optically-targeted methods^31^.

Despite providing a cautionary note for experimental designs using 4-OHT, our discovery provides an opportunity for new approaches to studying devaluation. 4-OHT can profoundly alter behavior after a single dose. In the assays we tested, this effect is more powerful than commonly used compounds like LiCl. Indeed, 4-OHT-induced devaluation even occurs when paired with alcohol indirectly, as water was also present in the homecage in our 2BC assays. The use of 4-OHT to drive activity-dependent recombination allows for the tagging of neurons that mediate these powerful effects.

The 4-OHT-mediated alcohol devaluation procedure has considerable experimental advantages over other similar assays such as alcohol adulteration with quinine^20,21^. Our conditioning procedure only changes the relationship between the organism and the stimulus rather than changing the stimulus itself, indicating that a mechanism that does not involve changing the sensory representation of the stimulus itself can devalue a stimulus. As a first step in the search for this type of mechanism, we combined TTP-S with graph theoretical analysis to show that activated brain areas with high PageRank scores are more tightly correlated with conditioned taste avoidance caused by 4-OHT delivery. Future experiments must uncover the precise functional role of these identified brain areas in mediating devaluation. Elucidating mechanisms of devaluation could contribute to the development of new treatments for addiction, where devaluing addictive compounds is clinically beneficial. While 4-OHT or related compounds may or may not be candidate treatments, understanding the mechanisms by which they mediate devaluation could prove critical in pinpointing novel targets in the brain for future therapeutics.

## Acknowledgements

The authors would like to thank Drs. Roberto Gulli, Rahim Hashim, George H. Denfield, Samantha A. Keil, Emily Parker, and Ioannis Koutlas for constructive comments on the manuscript and underlying data. Imaging was supported by the Zuckerman Institute’s Cellular Imaging platform.

## Author Contributions

EJK conceptualized the research ideas and methodology with input from CDS. EJK, RS, ME, SV, and LR ran the experiments, and EJK, RS, ME, SV, LR, and AR conducted the analyses. EJK created the visualizations and wrote the original draft of the manuscript. EJK and CDS revised the manuscript. CDS and EJK procured funding for the work, and CDS provided supervision of all research activities. All authors provided feedback and approved the final version of the manuscript.

## Methods

### Mice

C57BL/6J mice (stock #000664), homozygous Fos2A-iCreER/2A-iCreER (TRAP2) mice (stock #030323)^6^, and R26Ai14/+ (Ai14) mice (stock #007914)^32^ were obtained from Jackson Labs. TRAP2 mice were crossed in-house to Ai14 mice to obtain double heterozygous TRAP2:Ai14 mice as reported previously^5^. All mice were housed in a reverse light:dark cycle (12h:12h) room for at least 2 weeks before experimentation began. All behavioral experiments were conducted during the dark cycle. Unless otherwise noted, mice were allowed *ad libitum* access to food and water. Mice of both biological sexes were used in all experiments except for the 4-OHT dose response experiment (all females; see below). Mice were 8-16 weeks of age when experimentation began. Littermates were randomized to experimental or control groups, and all groups consisted of approximately equal numbers of males and females (maximum allowable difference being n=1 greater male or female per group). All procedures were performed in accordance with standard ethical guidelines and approved by Columbia University Institutional Animal Care and Use Committee.

### Drugs

4-hydroxytamoxifen (4-OHT; Sigma #H6278) was dissolved in 200 proof ethanol at a concentration of 20mg/mL and mixed thoroughly. The 4-OHT/EtOH mixture was added to a 1:4 mixture of castor oil:sunflower seed oil (Sigma #259853 and #S5007, respectively), and the ethanol was evaporated by vacuum centrifugation to arrive at a final concentration of 10mg/ml of 4-OHT. Mice were given i.p. injections of 4-OHT at doses of 50mg/kg, 25mg/kg, 10mg/kg, and 1mg/kg (4-OHT was further diluted in oil mixture for 10mg/kg and 1mg/kg doses for ease of dosing). We note that this oil mixture and vacuum centrifugation method for 4-OHT is the same that was reported in the initial Trap2 paper^6^ and has been widely used since. Other preparations, such as suspension in DMSO and Tween 80, have been used previously^33,34^, though the time-course of 4-OHT action appears similar between buffer preparations (maximum recombination at ∼2-3 hrs post-injection with minimal recombination after 4 hrs^6,33^). Castor oil:sunflower oil mixture was used as vehicle control for all 4-OHT experiments. Tamoxifen (Sigma #T5648) was dissolved in 200 proof ethanol at a concentration of 20mg/mL and mixed thoroughly. The tamoxifen/EtOH mixture was added to an equal amount of sunflower oil prior to evaporation of the ethanol by vacuum centrifugation. Additional sunflower oil was added to create working solutions prior to i.p. injections at 8mg/kg and 2mg/kg. Sunflower oil was used as vehicle control for tamoxifen experiments. Clozapine-N-oxide (CNO) dihydrochloride (HelloBio #HB6149) was dissolved in deionized water to create a 20mg/mL stock solution. On the day of CNO injection, stock solution was dissolved in 0.9% PBS to create a 5mg/mL working solution, and mice were given i.p. injections of 5mg/kg. PBS was used as vehicle control for CNO experiments. Lithium chloride (LiCl; Sigma #213233) was dissolved in deionized water to a concentration of 25mg/kg and injected at a dose of 125mg/kg, a dose commonly used to drive aversion and malaise (e.g.,^18^). PBS was used as vehicle control for LiCl experiments. All solutions were made fresh on the day of use.

### Two-bottle choice (2BC) assays

Mice were single housed and habituated to drinking tap water out of two glass bottles (Amuza, Inc.) in the homecage for at least two weeks. For 5d on / 2d off experiments, mice were given 24h access to 3% alcohol (v/v) for 2d in one bottle followed by 7% alcohol (v/v) for 3d. Mice then drank only water in both bottles during a 2d enforced withdrawal / washout period, as described previously^5^. 10% alcohol (v/v) was then introduced into one bottle for 5d, followed by another 2d washout period. This 5d / 2d cycle was repeated until the end of the experiment. Mice were given sham injections of saline for at least 4d prior to drug treatment (e.g., 4-OHT or tamoxifen). Drug injections were given 4h after 10% alcohol was placed into the cage (approximately 14:00, with alcohol placed into cages at lights off at 10:00) corresponding to the typical timing of maximal alcohol consumption^35^.

A modified intermittent access 2BC (IA2BC) assay was designed to explore the effects of 4-OHT on binge-like alcohol consumption. Single-housed mice were habituated to drinking water out of two bottles for at least two weeks. Mice were then exposed to increasing concentrations of alcohol in one bottle while alcohol remained in the other bottle, and the amount of time alcohol was available gradually decreased. Alcohol was only available 3d/week (Mondays, Wednesdays, and Fridays). On the first Monday of alcohol exposure, 3% alcohol (v/v) was available for 24h. On the first Wednesday, 3% alcohol was available for 6h. On the next two drinking days (Friday and the following Monday), 7% alcohol (v/v) was available for 5h and 4h, respectively. On the next two drinking days (a Wednesday and Friday), 10% alcohol (v/v) was available for 3h each day. Water was always available, even during alcohol access periods, for the quantification of alcohol preference compared to water. On each subsequent drinking day, mice were given 3h access to 20% alcohol (v/v). Mice were given sham injections of saline for at least 4d prior to drug treatment. On the 4-OHT treatment day, mice were injected with 4-OHT or vehicle 30 min after the first 20% drinking session. Mice continued to have access to 20% alcohol in one bottle for 3h each Monday, Wednesday, and Friday until the conclusion of the experiment.

A continuous access 2BC with a crossover design was designed to probe the effects of 4-OHT on alcohol preference while 1) alcohol was always present in the homecage, and 2) after mice had 4 weeks of experience drinking alcohol. Mice were single housed and habituated to drinking out of two bottles as described above. Mice were then exposed to 3% alcohol (v/v) in one bottle for 3d followed by 7% alcohol (v/v) in one bottle for 4d, with water always available in the other bottle. Mice were then given access to 10% alcohol (v/v) for the rest of the experiment, with 4-OHT or vehicle given 4h after 10% alcohol was introduced into one of the bottles. Following the normalization of the effect of 4-OHT on alcohol preference compared to vehicle-treated mice, we performed a crossover (after mice had been exposed to alcohol for 29 total days) whereby mice that previously received vehicle received 50mg/kg 4-OHT, and vice versa. Mice were given sham injections of saline for at least 4d prior to drug treatment.

For all 2BC experiments, bottles were weighed daily and the amount consumed recorded. Bottles were switched between sides every two days to mitigate side bias. Bottles of alcohol and water were kept in an empty cage for monitoring of evaporation and loss of fluid, though we did not correct for this given that approximately 90% of bottles in the empty cage lost between 0.0 and 0.1mL of fluid in 24h. Mice were weighed at least weekly for calculation of weight-corrected consumption.

### Blood alcohol quantification

Blood alcohol concentrations (BACs) were measured using an Analox AM1 (Analox Instruments) in n=16 mice after 5 weeks of drinking in the 5d on / 2d off 2BC. Blood was collected by severing the right atrium upon mouse sacrifice 4h into a 10% alcohol drinking session. Whole blood was centrifuged for plasma collection, and plasma was frozen at -80◦C until measurement. Quantification revealed an average BAC of 75.6 ± 9.0 (mean ± SEM) mg/dL in this assay, and BAC was statistically correlated to g/kg of alcohol consumed during the preceding 4 hours (Pearson r=0.76; *p*=0.0006; n=16).

BACs were similarly measured in the short access IA2BC assay. Blood was collected and processed as described above, with mice being sacrificed immediately after a 3h session where 20% alcohol was available following 5 weeks of alcohol consumption. Quantification revealed an average BAC of 123.0 ± 7.2 (mean ± SEM) mg/dL with all mice having BAC ≥ 100 mg/dL.

### Conditioned taste avoidance (CTA)

Mice were single housed and their water consumption was restricted for 20h prior to the first habituation day. On habituation days, a bottle filled with water and another empty bottle were placed in the homecage for 30 min. After this, water consumption was quantified, and mice were given a sham injection of 0.1mL PBS 30 min after the cessation of water consumption. On the conditioning day, mice were given access to an empty bottle and a bottle filled with a 0.06% grape or orange Kool-Aid (Kraft Heinz, Inc.) solution sweetened with 0.3% saccharin (Sigma #S1002) for 2 hours to overcome novelty-suppressed consumption. Mice were injected with either vehicle or the drug of interest 30 min after cessation of the novel flavor access session. Animals that consumed less than 0.3mL of the conditioned flavor were excluded from the experiment. Two hours after injections, mice were given access to *ad libitum* water in the homecage for at least 48 hours. Preference was then assessed via a 2BC task whereby animals had continuous access to both water and the conditioned Kool-Aid solution for 24-48 hours, with the location of bottles switched halfway through the access session and bottle weights recorded daily. During water restriction, animals were monitored to ensure they maintained at least 85% of their total body weight.

Similar to prior studies^18^, we assessed whether exposure to 4-OHT could lead to a CTA when mice were exposed to a familiar flavor. To do so, we habituated animals as described above, but in one group of mice (the ‘familiar’ group) we allowed access to the sweetened Kool-Aid solution for 2 conditioning sessions prior to drug exposure. Practically, this meant that mice in the familiar group were exposed to the Kool-Aid solution twice (paired with sham PBS injections 30 min post-consumption) prior to the conditioning day. Similar work has shown that LiCl is not effective in eliciting a CTA with a familiar flavor^18^.

### Conditioned place avoidance (CPA)

We utilized a custom-built conditioned place avoidance (CPA) behavior box with two compartments (approximately 10 cm x 20 cm floor area) separated by a central partition with a small opening to allow passage between compartments. One compartment had solid lines cut out of the acrylic floor and vertical black-and-white striping on the walls, while the other compartment had small holes cut into the floor and horizontal striping on the walls. The CPA apparatus was housed in a behavioral box lit by dim overhead light strips so that the center of each compartment measured ∼50-75 lux. Mice were given a 5 min habituation session where they were allowed to explore both compartments on the first day of the experiment. On the next day, mice were allowed to explore both compartments for 10 min (‘pre-test’ session) while being recorded from an overhead camera. Mice then underwent 3 conditioning sessions each for vehicle and 4-OHT, during which mice were confined to a single compartment for 45 min immediately following an injection of vehicle or 4-OHT (50mg/kg). Conditioning sides were randomly interleaved and unbiased. Mice were allowed 48hrs to recover following the last conditioning session. We then conducted a ‘post-test’ session in which mice were allowed to freely explore both sides of the CPA testing area for 10 min. An observer manually quantified the time spent (s) in each compartment in pre- and post-test sessions, which was also compared to automated quantification with DeepLabCut (see below).

### DeepLabCut behavioral analysis

CPA videos from pre- and post-test sessions were analyzed using DeepLabCut^36,37^. 20 frames were extracted from each of the 18 videos (n=9; pre- and post-test sessions) and the following body parts were labeled in each frame: snout, head center, left ear, right ear, neck center, left side, body center, right side, left hip, right hip, tail base, tail center, and tail tip; along with the following arena points: top left corner, top middle (at the position of the partition), top right corner, bottom right corner, bottom middle (at the position of the partition), and bottom left corner. We trained a ResNet-50 model on labeled data at 450,000 iterations and analyzed the full videos using this model. The analyzed and labeled videos were used to confirm proper identification of mouse body segments. The resulting coordinates were used to quantify mouse body position (body center label) and distance traveled in the pre- and post-test sessions.

An additional DeepLabCut model was trained on open field test (OFT) data described below. The same body points were labeled from each video as for the CPA analysis, as well as the four corners of the behavioral arena. We trained a ResNet-50 model on labeled data at 250,000 iterations and analyzed the full videos using this model. The analyzed and labeled videos were used to confirm proper identification of mouse body segments. The resulting coordinates were used to quantify mouse body position (body center label), including time spent in a 15-inch x 15-inch center zone, and distance traveled.

### Sucrose preference test (SPT)

Single housed mice were habituated to drinking water out of two bottles in the homecage for at least two weeks. In the sucrose ramp week, mice were exposed to 0.3% sucrose in one bottle (water in the other) for 2d, followed by 0.7% sucrose for 3d. Mice then drank only water in both bottles during a 2d enforced washout period as described above in the 5d on / 2d off alcohol 2BC experiments. Mice were then exposed to 1% sucrose in one bottle (water in the other) for 5d. On the first day of 1% sucrose exposure, mice were given an i.p. injection of either vehicle or 4-OHT (50mg/kg) 4h after sucrose was introduced into the cage. After 5d of 1% sucrose exposure, mice again underwent a 2d enforced washout period where they drank only water followed by 5 additional days of 1% sucrose exposure. As for the 2BC, bottles were weighed daily for quantification of consumption.

### Open field test (OFT)

Trap2:Ai14 mice were tested in the open field after injections of vehicle or 4-OHT. Animals were habituated to handling and injection stress (with 0.1mL PBS injections) for 3d prior to OFT testing. Animals were injected with either vehicle, 1mg/kg 4-OHT, or 50mg/kg 4-OHT 30 min prior to OFT testing. The OFT arena measured 24-inches x 24-inches with 12-inch-high walls, and the center zone was defined as a central square measuring 15-inches x 15-inches. Lighting was optimized so that the middle of the arena measured 400-500 lux. An overhead camera recorded animal behavior for 10 min in the open field. DeepLabCut was used to quantify mouse position and distance traveled, and we manually quantified behavioral epochs where the mouse was rearing, still and alert, or grooming (separated into paw grooms and full grooming sequences^38^).

### Chemogenetic activation of homecage ‘Trap’ ensemble

Trap2 mice were anesthetized with 4% vaporized isoflurane and given a retroorbital injection of either AAV-PHP.eb-hSyn-DIO-hM3D(Gq)-mCherry^19^ (Addgene #44361) or AAV-PHP.eb-hSyn-EGFP (Addgene #50465) at a titer of 2.0x10^11^ viral genomes per mouse diluted in 0.1mL of PBS. Mice were treated with ophthalmic ointment and monitored for signs of infection, and mice were single housed following the injection. After three weeks (to allow for viral penetration to the CNS), mice were given 50mg/kg 4-OHT in their homecages to ‘trap’ an ensemble unrelated to any specific stimulus. After four weeks, mice underwent a CTA as described above (5d habituation followed by 2d break with *ad libitum* access to water). On the conditioning day, mice were allowed to consume a sweetened Kool-Aid solution and given an injection of 5mg/kg CNO or vehicle 30 min after cessation of consumption.

*Quantification of AAV-PHP.eb-hSyn-DIO-hM3D(Gq)-mCherry expression using BrainJ*

Approximately two weeks after the cessation of the CTA experiment, Trap2 mice previously injected with AAV-PHP.eb-hSyn-DIO-hM3D(Gq)-mCherry or AAV-PHP.eb-hSyn-EGFP were anesthetized with a ketamine/xylaxine (100mg/kg and 10mg/kg, respectively) mixture prior to transcardial perfusion with 20mL of PBS followed by 20mL of 4% paraformaldehyde (PFA), as described previously^5^. Brains were extracted and left overnight in a 4% PFA solution before being transferred to PBS. Brains were cut into 50-micron sections on a vibratome (Leica, Inc.) and washed three times in PBS prior to mounting. After sections were coverslipped and allowed to dry, confocal images of whole sections were obtained using a Yokogawa W1 spinning disk confocal microscope using a 4x 0.2 NA objective tiled to cover the whole section in the 405nm and 561nm channels.

Images were processed using the BrainJ^30^ plugin for FIJI^39^. Images corresponding to each 50-micron section through the brain were processed to number the sections sequentially from anterior to posterior and remove background to be compatible with the BrainJ whole-brain registration. Images are centered and rotated to facilitate registration for each individual brain section, which yields a 3D brain volume rendering that is then registered to the Allen Brain Atlas Common Coordinate Framework (CCF v3) using elastix^40^. As described previously^5^, registrations were examined for proper alignment, and all n=6 brains from the AAV-PHP.eb-hSyn-DIO-hM3D(Gq)-mCherry + CNO CTA experiment were adequately registered. In order to quantify the number of neurons in AAV-PHP.eb-hSyn-DIO-hM3D(Gq)-mCherry brains, we trained an Ilastik^29^ model within BrainJ to identify transfected cells, projections, and background images prior to segmentation. The model was trained on three images from each of the n=6 brains for a total of 18 training images. Model quality was assessed in real-time using the Ilastik GUI. We then used BrainJ to count mCherry-positive cells across the entire imaged portion of the brain. Projections were quantified for quality control. An average of 274,641 (SD = 87,506; range = 183,439 - 392,742) mCherry-positive neurons were detected in each brain, remarkably similar to the average number of tdTomato-positive neurons detected in whole-brain studies of stimulus-evoked ensembles in Trap2:Ai14 mice (average of 239,680 neurons)^5^. Example training and quantification images are shown in Extended Data Fig. 8.

### Brainwide activity-dependent labeling with TTP-S

We utilized a recently developed activity-dependent screening pipeline called two-timepoint statistical inference with subtraction (TTP-S)^5^. TTP-S relies on neuronal activity labeling at two independent timepoints, such that the overlap in activity (experimentally determined by double-labeling of cells in Trap2:Ai14 mice) can be compared to chance levels of overlap using a hypergeometric distribution. Mice are given 50mg/kg 4-OHT at timepoint 1 (TP1) to indelibly label active neurons with tdTomato, followed by sacrifice for whole-brain processing and cFos labeling at timepoint 2 (TP2).

Trap2:Ai14 mice underwent a 5d on / 2d off alcohol 2BC assay as described above. For the experimental group, which we call ethanol-ethanol or EE mice, TP1 occurred on the first day of exposure to 10% alcohol in one bottle approximately 4 hours after 10% alcohol was introduced into the homecage. TP2 occurred one week later approximately 4 hours after re-exposure to 10% alcohol in one bottle in the homecage following the 2d enforced withdrawal / water-only washout period. TTP-S utilizes a ‘subtraction’ group of mice, whereby mice are exposed to two distinct stimuli at TP1 and TP2. Therefore, any double-labeled cells likely represent non-specific neural activity (e.g., tonically active neural populations, neurons responding to non-stimulus environmental features)^5^. For the subtraction group, which we call ethanol-homecage (water) or EH mice, TP1 was identical to the EE group, and TP2 occurred at the same time (one week after TP1) but without exposure to any alcohol. We also generated groups of mice where TP1 and TP2 occurred in the homecage without any specific stimulus exposure (homecage-homecage or HH mice) and where TP1 occurred after seven total weeks of alcohol consumption in the 5d on / 2d off 2BC assay and TP2 occurred one week later approximately 4hrs after re-exposure to 10% alcohol (late-late or LL group).

Mice were anesthetized and perfused as described above. Brains were removed and stored in 4% PFA overnight prior to transfer to PBS for 24 hrs followed by 20% sucrose in PBS for 48hrs. Brains were then embedded in Tissue-Tek Optimum Cutting Temperature (OCT; Sakura Finetek, #4583) medium prior to flash freezing and storage at -80°C until sectioning. As described previously^5^, brains were sectioned on a cryostat (Leica) at 50-microns through the entire brain, and floating sections were collected in PBS. For immunostaining, sections were washed three times in PBS and incubated in 5% donkey serum (Jackson Immunoresearch; #017-000-121) in PBS with Triton-X (PBST) for 1hr. Sections were stained with primary antibodies targeting c-Fos (Cell Signaling, #2250, 1:1000 dilution in PBST and 5% donkey serum) and NeuN (EMD Millipore, #MAB377, 1:1000) and incubated at 4°C overnight. Sections were then washed three times with PBST prior to conjugation with secondary antibodies: AlexaFluor-647 (Invitrogen #A31573, 1:1000) and AlexaFluor-488 (Life Technologies #A21202, 1:1000), as well as a DAPI counterstain (Sigma-Aldrich, #D9542 1:5000). Sections were mounted and coverslipped with mounting medium. Confocal images were obtained using a Yokogawa W1 spinning disk confocal microscope using a 4x 0.2 NA objective in 4 channels: 405nm (DAPI stain), 488nm (anti-NeuN stain), 561nm (tdTomato label – TP1), and 647nm (anti-c-Fos stain – TP2).

Sections were reordered and registered to the Allen Brain Atlas CCF v3 as described previously^5,30^ and in an identical manner to AAV-PHP.eb-hSyn-DIO-hM3D(Gq)-mCherry brains described above. Only brains that could be properly registered to the CCF were used for further processing and analysis (n=5 for EE and EH groups; n=4 for HH group; n=6 for LL group). We utilized an updated version of BrainJ^5^ to quantify cells in the DAPI, tdTomato, and anti-cFos channels. This version of BrainJ automatically uses CARE/RCAN networks and custom U-Nets to segment and detect cells in each channel, and we considered cells double-labeled by using an algorithm that combines information on the distance between centroids and fraction of overlapping volume between two cells. Models and BrainJ-Python code are available at https://github.com/lahammond/brainj-py.

### Systematic review of effects of re-activating or inhibiting a previously ‘trapped’ ensemble

We conducted a broad search of the literature for studies utilizing Trap2 or FosTrap mice in PubMed and Web of Science using the following search strategy; PubMed: ("TRAP2" OR "FosTRAP" OR "targeted recombination in active populations") AND ("mouse" OR "mice"); Web of Science: (ALL=(("mouse" OR "mice"))) AND ALL=(("TRAP2" OR "FosTRAP" OR "targeted recombination in active populations"))). The authors added literature from their own collection of papers that were relevant to the query. Only papers published between within the preceding decade (2017 onward) were included. Queries were last performed on March 27^th^, 2026. The results were reviewed in Covidence (Veritas Health Information) and duplicates were removed. Studies were evaluated following inclusion criteria: use of Trap2 (or Trap2:reporter line) or FosTrap mice, use of 4-OHT for ‘trap’ procedure, activation or inhibition (via chemogenetics or optogenetics) of previously ‘trapped’ ensemble, and activation/inhibition of this ensemble in a behavioral task that could be interpreted in a straightforward manner as aversive (or anxiogenic) or appetitive (or anxiolytic). These included CPP/CPA, CTA, OFT, elevated plus/zero maze, and fear conditioning. Outcomes were classified as aversive or appetitive, and the quality of each study was additionally recorded.

### Statistical analysis

#### Behavioral assays

2BC and SPT assays were analyzed with two-way repeated measures (RM) ANOVA (factors: group, time, group x time interaction). Analyses were split into pre-treatment and post-treatment periods. In order to control for day-to-day variation in preference and consumption, *post-hoc* testing was performed on the weekly average of alcohol preference and consumption metrics using Šidák’s tests correcting for multiple comparisons. We note that this choice was deliberately conservative compared to other methods for analyzing 2BC drinking data, such as performing repeated Fisher’s least significant difference tests or uncorrected Tukey tests on individual daily values. For instance, there are daily alcohol preference values that show significant alterations between groups (lower alcohol preference in 4-OHT-treated mice compared to vehicle) in the SPT shown in Fig. 2 and the continuous alcohol access 2BC shown in Extended Data Fig. 3. Given this likely inflation of type II error, we have taken caution to not overinterpret the biological and behavioral significance of these results and emphasizing only that single doses of 4-OHT and tamoxifen appear to change behavior in a temporally extended manner. The alternative, in which we would correct for multiple comparisons across all singular days (or timepoints) post-injection, is also not ideal given that it would, at times, necessitate correction for >30 comparisons and thus inflate type I error. Note that we only considered omnibus significance of tests of group and group x time as sufficient to advance to *post-hoc* testing, as effects of time were expected in consumption experiments with ramping concentrations of alcohol and sucrose. SPT data were analyzed in an identical manner (two-way RM ANOVA with *post-hoc* multiplicity-corrected Šidák’s tests on weekly average values). For the 4-OHT dose response group, week two (after 4-OHT exposure; given the results from female C57BL/6J mice showing decreased alcohol preference for only ∼5 days) was tested using a one-way ANOVA followed by *post-hoc* Holm-Šidák tests correcting for multiple comparisons. For Trap2:Ai14 mice shown in Extended Data Fig. 1, we used a mixed effects models (restricted maximum likelihood; REML, with *post-hoc* multiplicity-corrected Šidák’s tests on weekly average values) with factors of group, time, and group x time interaction as some mice in this experiment were sacrificed for tissue collection during the behavioral assay. Alcohol preference was calculated per mouse per day as:

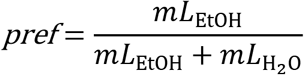

where *pref* is alcohol preference and *mL*_EtOH_ and *mL*_H2O_ are the amount of alcohol and water consumed (in mL), respectively. Unless otherwise specified (e.g., for the modified IA2BC assay), only alcohol consumption days were included in analyses.

For CTA data, flavor preference was calculated similarly to alcohol preference. Preference values were compared with unpaired *t*-tests (for two-group data), Kruskal-Wallis ANOVA with *post-hoc* Dunn’s test (for three-group data not normally distributed), and one-way ANOVA with post-hoc Tukey tests (for normally distributed three-group data). For CPP data, time spent in each compartment was compared for pre- and post-test timepoints using unpaired *t*-tests both for manually quantified and DeepLabCut data. Distance traveled (cm; from DeepLabCut) was compared between pre- and post-test timepoints with a paired *t*-test. We compared manual and DeepLabCut analyses by performing Pearson’s correlation and simple linear regression on values for time spent in each compartment. For OFT time, time spent in the center and distance traveled (from DeepLabCut) were analyzed with one-way ANOVAs, and manually quantified behaviors (still and alert; rearing; grooming) were analyzed with a two-way ANOVA.

#### Brainwide correlations with preference for conditioned stimuli

Per-region counts of mCherry-positive cells (from n=6 AAV-PHP.eb-hSyn-DIO-hM3D(Gq)-mCherry injected Trap2 mice treated with CNO during CTA described above) were combined between left and right hemispheres. Total mCherry-positive counts in the entire brain were examined for a correlation with CTA flavor preference using Spearman’s correlation (with Pearson also performed given small sample size). Spearman was used for per-region correlation of mCherry-positive cells with CTA flavor preference. We performed Spearman correlations for each brain region corrected for region size^41^. A volcano plot was used for visualization, and 12 brain regions displayed a *p*-value below 0.05 but no region met Benjamini–Hochberg false discovery lab (FDR) criteria.

We used brain regions that met significance criteria from our TTP-S analysis of alcohol devaluation (see below) to further parse the brainwide data examining the correlation between regional AAV-PHP.eb-hSyn-DIO-hM3D(Gq)-mCherry positivity and CTA flavor preference. n=144 brain regions were 1) significant in the EE-EH TTP-S analysis; and 2) demonstrated sufficient correlations with other brain regions (threshold of *r* = ±0.6). We performed Spearman correlations (as this data was not normally distributed) between the positive network PageRank values (see below) and the brain region correlation of mCherry-positive cells with CTA flavor preference to determine if hub brain regions with higher PageRank values might contribute differentially to behavior. The same analysis was done with negative network PageRank values. We plotted the Spearman correlation values between mCherry-positivity and CTA flavor preference for the top 20% of regions by PageRank (n=29) in both the negative and positive networks compared to the non-hub (n=115) regions. We created a null distribution by shuffling Spearman *r* values of the n=144 brain regions 100,000 times without replacement and testing the observed *r* value of the hub regions (by PageRank) with a two-sided permutation test corrected for multiple comparisons.

#### TTP-S statistical analysis and graph theoretical analysis

As described previously^5^, BrainJ-Python cell counts were combined across hemispheres and a probability distribution for dispersion of observed overlap (double-labeled cells at both TP1 and TP2) was calculated for each individual mouse brain:

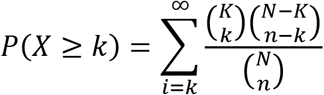

where *X* denotes the random variable giving the size of the overlap, *N* is the total cell count (from DAPI channel cell counts), *K* is the number of active cells at TP1, *n* is the number of labeled cells at TP2, and *k* is the number of double-labeled cells. The 1-sided *p*-value gives the probability that at least *k* neurons are double-labeled given by the sum of all probabilities either equal to or greater than *k*.

In our initial paper describing the TTP-S approach, we noted that using the above statistical test tends to yield many brain regions with higher overlap than expected by chance, likely representing false positives^5^. Thus, the subtraction portion of the TTP-S pipeline creates a null distribution for statistical testing across groups of mice by bootstrapping samples from an experimental group mouse (i.e., an EE mouse) and subtracting bootstrapped samples from a subtraction group mouse (i.e., an EH mouse). This is done for every pair of hypergeometric distributions from the two groups of mice. The null distribution (k_diff_) is created by performing the bootstrapping procedure sampling 90000 times with replacement from each distribution and then creating a new null distribution of the differences. *p*-values are computed with the new null distribution by summating the values in that distribution that are equal to or greater than the actual observed difference. *P*-values were computed for each pair-wise comparison of mice (i.e., n=5 in each group yields 25 pairwise comparisons) and combined using a harmonic mean approach^5,42^. Brainwide significance thresholds were set using a stringent Bonferroni correction (α=0.05/620), with n=175 brain regions meeting criteria for significance in the alcohol devaluation (EE-EH) analysis.

We used graph theoretical analysis to determine possible hub regions mediating the alcohol devaluation effect^5,43^ using the Brain Connectivity Toolbox in MATLAB^44^. We computed a Pearson correlation matrix consisting of normalized double-labeled cell counts (overlap / tdTomato) across mice from the significant brain regions identified from the EE-EH TTP-S analysis. Positive correlation matrices were thresholded at *r* > 0.6 and negative correlation matrices were thresholded at *r* < -0.6. Clusters were determined by performing the Louvain community detection algorithm on the positive correlation matrix 100 times and thresholding at a consensus of 0.5, though all modules reached full (100/100; 100%) consensus. Module enrichment for the major divisions of the CCF v3 were tested using Fisher’s exact tests not corrected for multiple comparisons using all regions input into the graph (n=144) and all brain regions counted. We calculated PageRank as well as participation coefficient (PC), within-module degree Z-score (WMDz), degree centrality, eigenvector centrality, and betweenness score for the positive and negative correlation matrix using the Brain Connectivity Toolbox. Nodes and edges were visualized in Gephi^45^ for the positive and negative networks independently using the Fruchterman-Reingold algorithm with no overlap allowed.

We examined the relationship between neural activity between groups (EE vs EH, EE vs HH, and EE vs LL) by comparing Pearson correlation matrices. Region x overlap-count matrices were created for each group excluding any regions with zero or NaN counts. To assess overall structural similarity between matrices, we computed the Spearman correlation between the upper triangles of the two matrices as an observed similarity statistic (Mantel test). A null distribution was generated by randomly permuting row and column indices of the target (non-EE) matrix across 5,000 iterations and recomputing the Spearman correlation with the EE matrix at each iteration. A one-tailed *p*-value was computed as the proportion of permuted Spearman values less than the observed value, testing whether the two matrices were more dissimilar than expected by chance. We also confirmed that brainwide activation patterns were best explained by group-level differences with PERMANOVA followed by PERMDISP testing^46,47^. Each mouse was represented as a vector of normalized double-labeled cell counts from all registered brain regions valid across all brains (n=456) that contained no zero or NaN counts. PERMANOVA assessed whether group membership explains a significant proportion of variance in Euclidian distance vector space using a *pseudoF* statistic computed from the distance matrix and tested against a null distribution generated by randomly permuting group labels over 5000 iterations (PERMANOVA *pseudoF*=1.7233; *p*=0.0012). PERMDISP then measured distances from each animal to its group centroid in coordinate space that were also tested via permutation (5000 iterations; PERMDISP *pseudoF*=2.2972; *p*=0.2904). A significant PERMANOVA with a non-significant PERMDISP supports the interpretation that group differences reflect centroid separation (group-level mean differences) without significant differences in dispersion between the groups. Custom MATLAB code described in detail previously^5^ and available at https://github.com/ekyzar/ttps_pipeline was used for all TTP-S and graph theoretical analysis.

Significance was set at α=0.05 and corrected for multiple comparisons unless otherwise specified. Detailed information on statistical test results can be found in Tables S5 and S6.

