## Extended Data Figures for "Lasting aversive consequences of a single dose of 4-hydroxytamoxifen"

Extended Data Figures 1-10.

Supplemental Tables S1-S6 – see additional document.

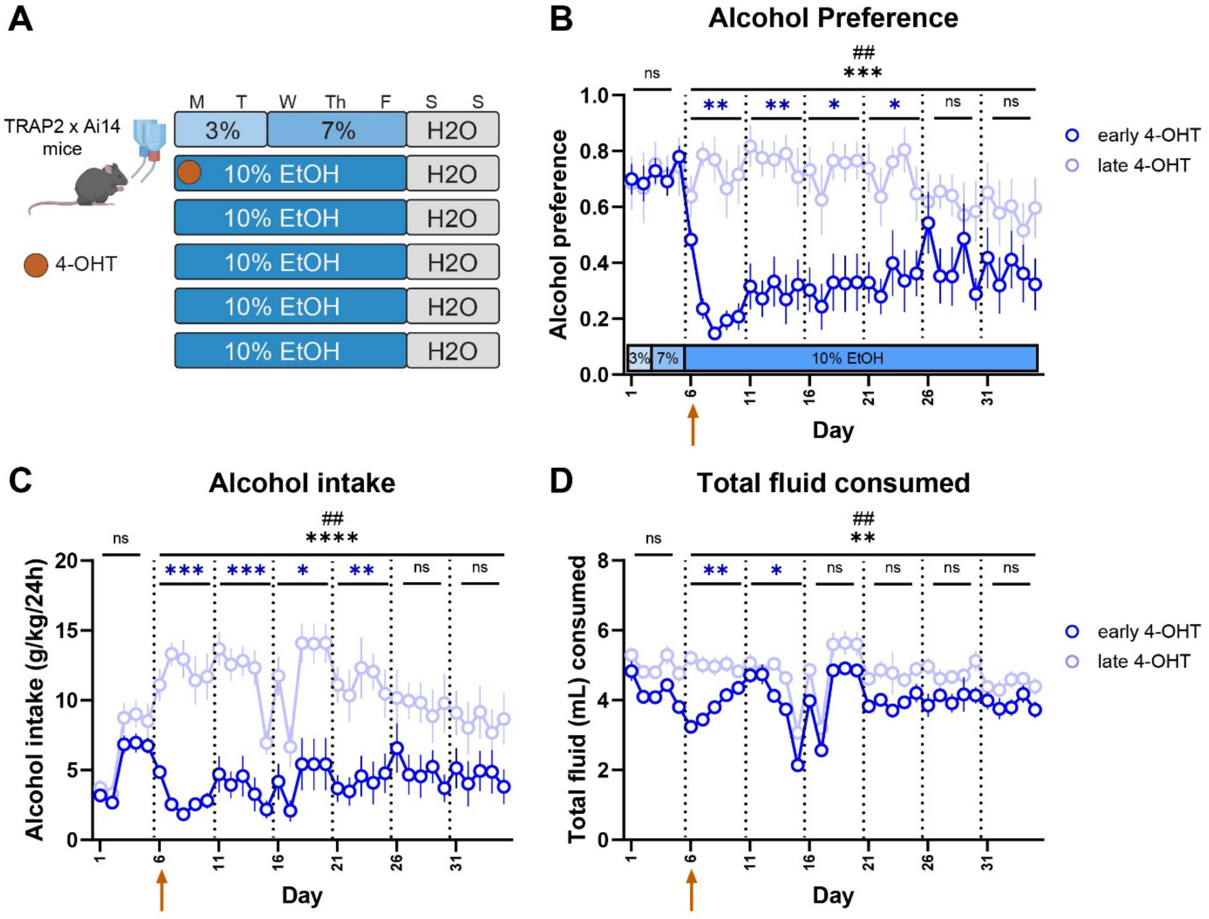


**Extended Data Figure 1. Alcohol preference and consumption in Trap2:Ai14 mice after a single dose of 4-hydroxytamoxifen (4-OHT). A)** Experimental schematic of a two-timepoint design comparing Trap2:Ai14 mice drinking alcohol in a two-bottle choice (2BC) task. Some mice were given 4-OHT (50mg/kg, i.p.) on the first day of 10% alcohol exposure (early 4-OHT group) while other mice were not given 4-OHT until >6 weeks of drinking experience (late 4-OHT group; see Methods). **B)** Alcohol preference (calculated as mL of alcohol consumed per day divided by total fluid consumed per day) in mice given 4-OHT (50mg/kg) on the first day of 10% alcohol exposure in a two-bottle choice (2BC) task. Arrow indicates day of 4-OHT treatment in early 4-OHT group. Mixed effects REML omnibus effects: ***$F_{Group}(1,22) = 21.91$, $p$=0.0001; $^{##}F_{GroupxTime}(6.116,92.38) = 3.580$, $p$=0.003. n=16 in 'early 4-OHT' group for first two weeks, n=8 in subsequent weeks; n=8 in 'late 4-OHT' group. Blue asterisks represent differences between early and late groups in the weekly rolling average of alcohol preference by multiplicity-corrected Šidák tests after mixed effects modeling. **$p$<0.01, *$p$<0.05. No significant effects in the pre-treatment period. **C)** Alcohol intake (g/kg/24h) in mice given 4-OHT (50mg/kg) on the first day of 10% alcohol exposure in a two-bottle choice (2BC) task. Arrow indicates day of 4-OHT treatment in early 4-OHT group. Mixed effects REML omnibus effects: ****$F_{Group}(1,22) = 30.06$, $p<0.0001$; ##$F_{GroupxTime}(5.319,80.34) = 4.375$, $p=0.0011$. n=16 in 'early 4-OHT' group for first two weeks, n=8 in subsequent weeks; n=8 in 'late 4-OHT' group. Blue asterisks represent differences between early and late groups in the weekly rolling average of alcohol intake by multiplicity-corrected Šidák tests after mixed effects modeling. ***$p<0.001$, **$p<0.01$, *$p<0.05$. No significant effects in the pre-treatment period. **D)** Total fluid intake (mL/day) in mice given 4-OHT (50mg/kg) on the first day of 10% alcohol exposure in a two-bottle choice (2BC) task. Arrow indicates day of 4-OHT treatment in early 4-OHT group. Mixed effects REML omnibus effects: **$F_{Group}(1,22) = 10.60$, $p=0.004$; ##$F_{GroupxTime}(6.803,102.8) = 3.105$, $p=0.006$. n=16 in 'early 4-OHT' group for first two weeks, n=8 in subsequent weeks; n=8 in 'late 4-OHT' group. Blue asterisks represent differences between early and late groups in the weekly rolling average of alcohol intake by multiplicity-corrected Šidák tests after mixed effects modeling. **$p<0.01$, *$p<0.05$. No significant effects in the pre-treatment period. All data shown as mean ± SEM.

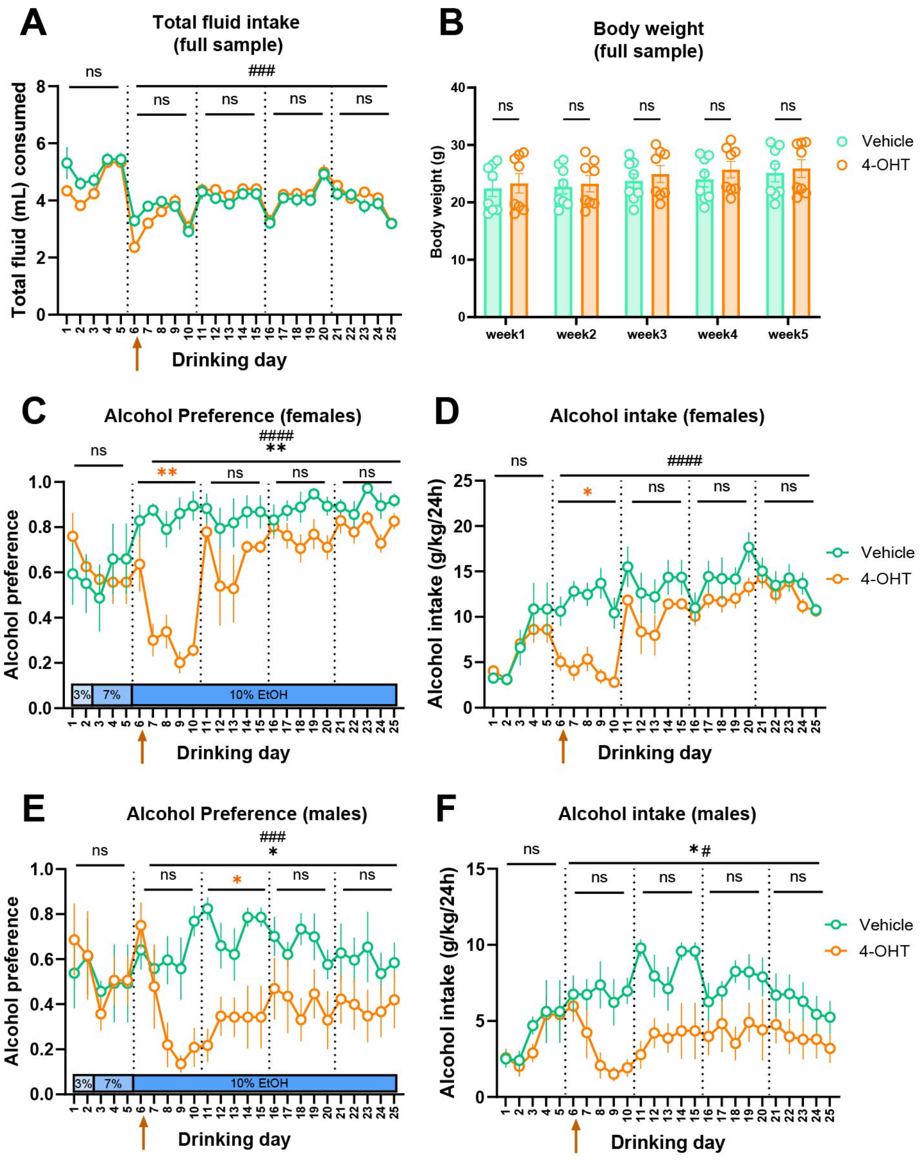


**Extended Data Figure 2. Alcohol consumption and preference in C57BL/6J mice after a single dose of 4-hydroxytamoxifen (4-OHT). A)** Total fluid intake (mL/day) for each mouse in full sample (both males and females). Arrow indicates day of 4-OHT treatment in early 4-OHT group. 2-way RM-ANOVA omnibus effects: $F_{Group}(1,14) = 0.066$, $p$=0.80; $^{###}F_{GroupxTime}(19,266) = 2.809$, $p$=0.0001. There were no significant differences in the weekly average total fluid intake between the groups by multiplicity-corrected Šidák's tests. No significant effects in pre-treatment period. n=8. **B)** Body weight (kg) in full sample (both males and females) measured weekly. No significant effects by 2-way RM ANOVA; n=8. **C)** Alcohol preference in females in 2BC task. 2-way RM-ANOVA omnibus effects: $^{**}F_{Group}(1,6) = 16.51$, $p$=0.007; $^{####}F_{GroupxTime}(19,114) = 5.622$, $p$<0.0001. Orange asterisks represent differences between groups in the weekly rolling average of alcohol preference by multiplicity-corrected Šidák tests after 2-way RM ANOVA. **$p$<0.01. No significant effects in the pre-treatment period. n=4. **D)** Alcohol intake (g/kg/24h) in females. $F_{Group}$ (1,6) = 5.450; p=0.058; ####$F_{GroupxTime}$(19,114) = 4.973, $p$<0.0001. Orange asterisks represent differences between groups in the weekly rolling average of alcohol intake by multiplicity-corrected Šidák tests after 2-way RM ANOVA. *$p$<0.05. No significant effects in the pre-treatment period. n=4. **E)** Alcohol preference in males in 2BC task. 2-way RM-ANOVA omnibus effects: *$F_{Group}$(1,6) = 6.394, $p$=0.049; ###$F_{GroupxTime}$(19,114) = 2.773, $p$=0.0004. Orange asterisks represent differences between groups in the weekly rolling average of alcohol preference by multiplicity-corrected Šidák tests after 2-way RM ANOVA. *$p$<0.05. No significant effects in the pre-treatment period. n=4. **F)** Alcohol intake (g/kg/24h) in males. *$F_{Group}$ (1,6) = 6.912; p=0.039; #$F_{GroupxTime}$(19,114) = 1.952, $p$=0.016. No significant effects between groups in the weekly rolling average of alcohol intake by multiplicity-corrected Šidák tests after 2-way RM ANOVA. No significant effects in the pre-treatment period. n=4. All data shown as mean ± SEM. Note that combined data (both males & females) shows significantly decreased alcohol preference and intake (see Fig. 1B,C).

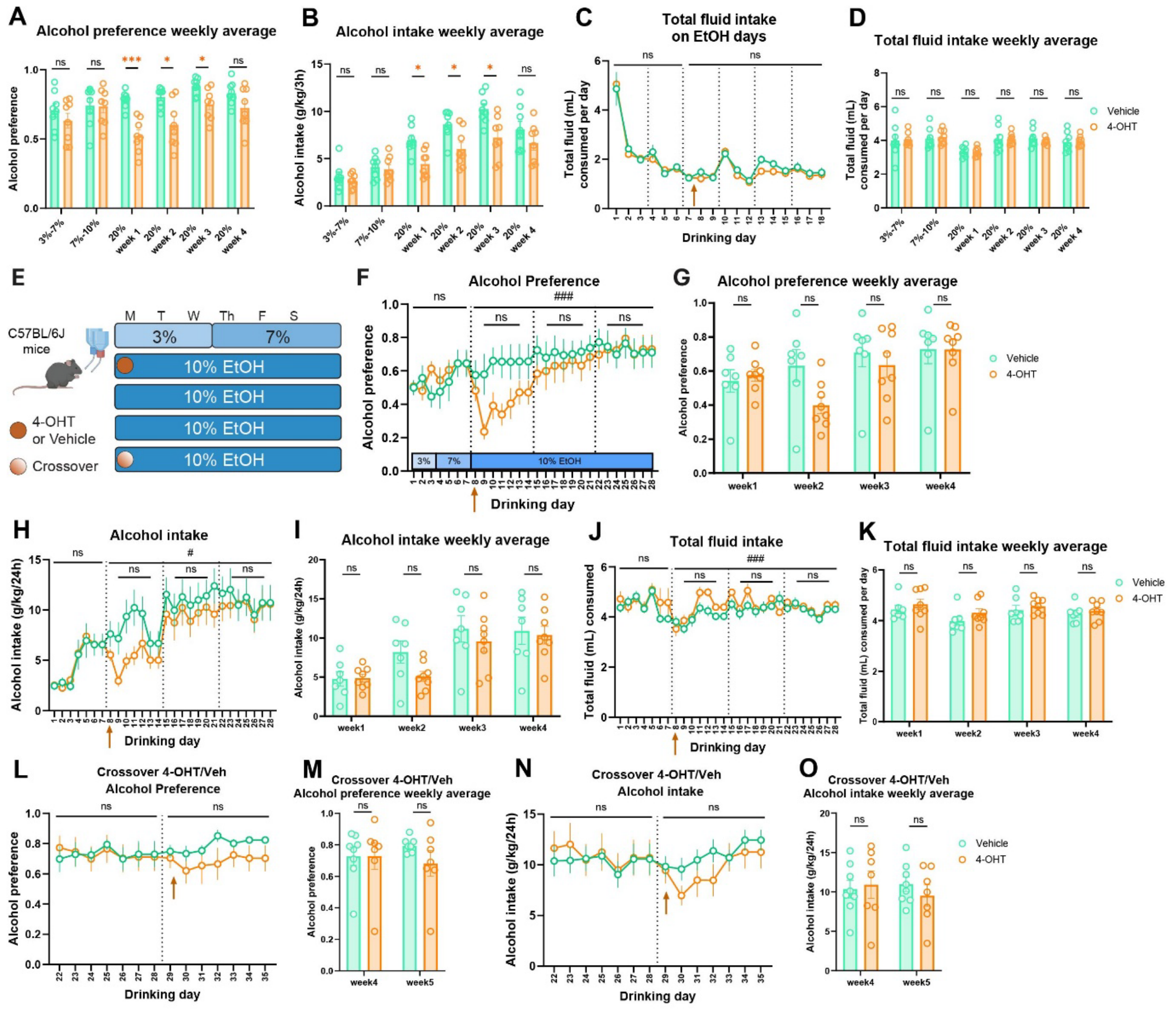


**Extended Data Figure 3. Alcohol consumption and preference in a short access binge consumption 2BC assay (A-D) and a continuous access 2BC with late crossover (E-O) in C57BL/6J mice after a single dose of 4-hydroxytamoxifen (4-OHT)**. **A)** Weekly average alcohol preference for visualization of individual values in each mouse in full sample (both males and females) of C57BL/6J mice after 4-OHT (50mg/kg; i.p.) on the first day of 20% alcohol exposure in a modified intermittent access two-bottle choice (2BC) assay. Orange asterisks represent differences between groups in the weekly rolling average of alcohol preference by multiplicity-corrected Šidák tests after 2-way RM ANOVA. ****p*<0.001, **p*<0.05. n=8. **B)** Weekly average alcohol intake (g/kg/24h) for visualization of individual values in each mouse in full sample (both males and females) of C57BL/6J mice after 4-OHT (50mg/kg) in modified intermittent access 2BC assay. Orange asterisks represent differences between groups in the weekly rolling average of alcohol intake by multiplicity-corrected Šidák tests after 2-way RM ANOVA. **p*<0.05. n=8. **C)** Total fluid intake (mL/day) in full sample (both males and females) of mice given 4-OHT (50mg/kg) on the first day of 20% alcohol exposure in a modified intermittent access 2BC task. Graph shows only fluid intake during short access alcohol consumption periods (see Methods). Arrow indicates day of 4-OHT treatment in early 4-OHT group. No significant effects by 2-way RM-ANOVA. n=8. **D)** Weekly average total fluid intake (mL/day) for each mouse in full sample (both males and females) for short access 2BC, including non-alcohol consumption days. No significant effects by multiplicity-corrected Šidák tests. n=8. **E)** Experimental schematic of exposure of C57BL6/J mice to 4-OHT (50mg/kg; i.p.) or vehicle on the first day of fully continuous 10% alcohol exposure with a crossover component. Mice exposed to vehicle on drinking day 8 were exposed to 4-OHT on drinking day 29. **F)** Alcohol preference in full sample (both males and females) of mice given 4-OHT on the first day of 10% alcohol exposure in a continuous access 2BC before crossover. $F_{Group}(1,13) = 1.106$, $p=0.31$; $^{\#\#\#}F_{GroupxTime}(20,260) = 2.425$, $p=0.0008$; No significant effects on weekly (7d) rolling average of alcohol preference by multiplicity-corrected Šidák tests (though see Methods for note on conservative approach to *post-hoc* multiplicity corrected significance testing). No significant effects in pre-treatment period. n=7-8. **G)** Weekly average alcohol preference for visualization of individual values for each mouse in full sample (both males and females) for continuous access 2BC. No significant effects by multiplicity-corrected Šidák tests; n=7-8. **H)** Alcohol intake (g/kg/24h) in full sample (both males and females) of mice given 4-OHT on the first day of 10% alcohol exposure in a continuous access 2BC before crossover. $F_{Group}(1,13) = 0.9318$, $p=0.35$; $^{\#}F_{GroupxTime}(20,260) = 1.738$, $p=0.028$; No significant effects on weekly (7d) rolling average of alcohol preference by multiplicity-corrected Šidák tests. No significant effects in pre-treatment period. n=7-8. **I)** Weekly average alcohol intake (g/kg/24h) for each mouse in full sample (both males and females) for continuous access 2BC. No significant effects by multiplicity-corrected Šidák tests; n=7-8. **J)** Total fluid intake (mL/day) in full sample (both males and females) of mice given 4-OHT on the first day of 10% alcohol exposure in a continuous access 2BC before crossover. $F_{Group}(1,13) = 1.013$, $p=0.33$; $^{\#\#\#}F_{GroupxTime}(20,260) = 2.410$, $p=0.0009$; No significant effects on weekly (7d) rolling average of alcohol preference by multiplicity-corrected Šidák tests. No significant effects in pre-treatment period. n=7-8. **K)** Weekly average total fluid intake (mL/day) for each mouse in full sample (both males and females) for continuous access 2BC. No significant effects by multiplicity-corrected

Šidák tests; n=7-8. **L)** Alcohol preference in full sample (both males and females) in continuous access 2BC after crossover, where mice given 4-OHT previously received vehicle (as these mice had returned to baseline levels of alcohol preference and consumption) and mice previously given vehicle received 4-OHT. No significant effects by 2-way RM ANOVA.n=7-8. **M)** Weekly average alcohol preference for each mouse in full sample (both males and females) in continuous access 2BC after crossover. No significant effects by multiplicity-corrected Šidák tests; n=7-8. **N)** Alcohol intake (g/kg/24h) in full sample (both males and females) in continuous access 2BC after crossover. No significant effects by 2-way RM ANOVA. n=7-8. **O)** Weekly average alcohol intake (g/kg/24h) for each mouse in full sample (both males and females) in continuous access 2BC after crossover. No significant effects by multiplicity-corrected Šidák tests; n=7-8. All data shown as mean ± SEM.

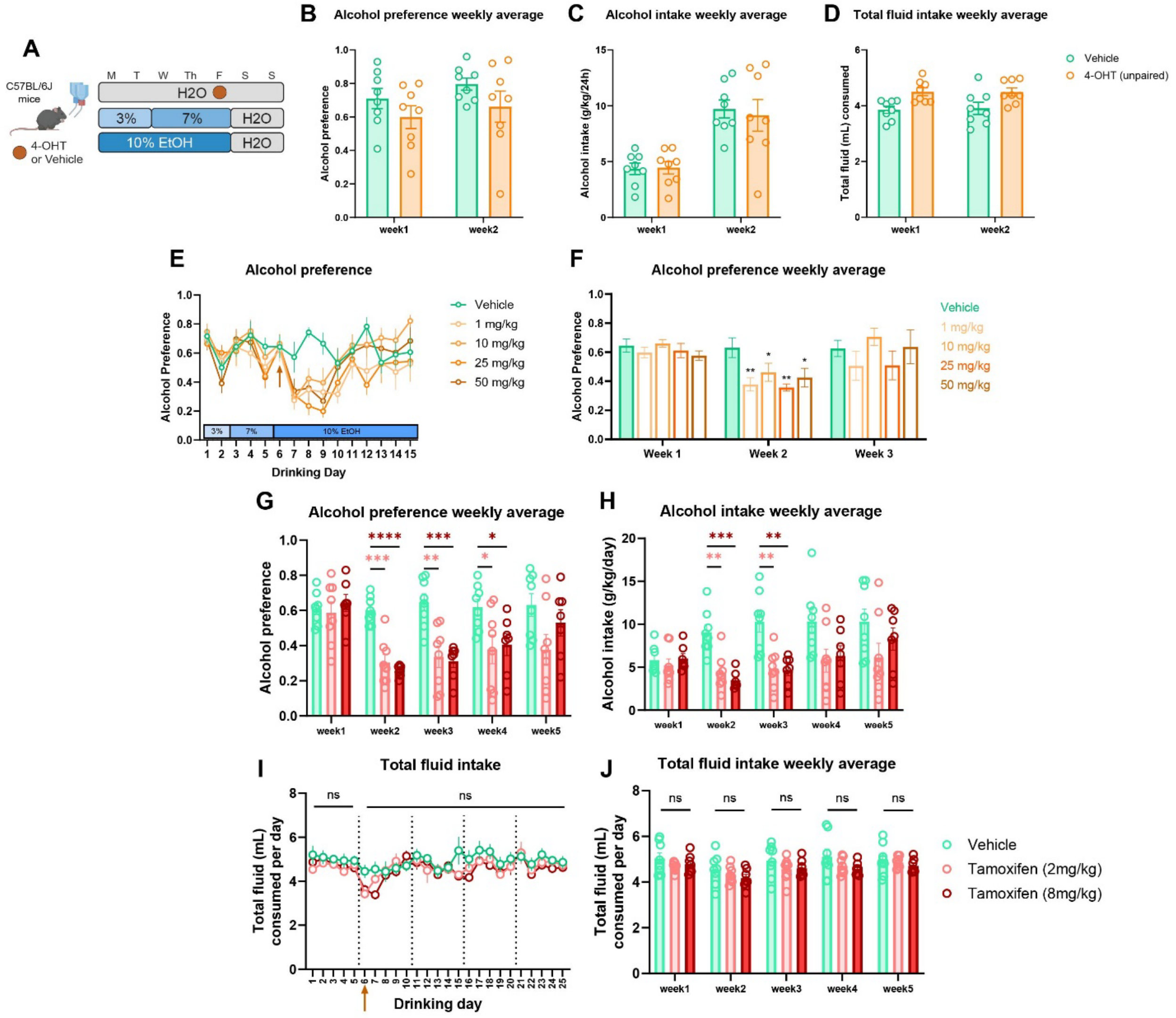


**Extended Data Figure 4. Alcohol consumption and preference in an unpaired two-bottle choice (2BC) assay with 4-hydroxytamoxifen (4-OHT), 4-OHT dose response 2BC, and tamoxifen 2BC in C57BL/6J mice**. **A)** Experimental schematic of C57BL/6J mice (n=8; 4 each males/females) given a single dose of 50mg/kg of 4-OHT i.p. not paired with alcohol prior to the introduction of escalating alcohol concentrations. **B)** Weekly average alcohol preference each mouse in full sample (both males and females) after unpaired 4-OHT. There were no significant differences in the weekly average of preference between the groups by multiplicity-corrected Šidák's tests. n=8. **C)** Weekly average alcohol intake (g/kg/24h) for each mouse in full sample (both males and females) after unpaired 4-OHT. There were no significant differences in the weekly average of intake between the groups by multiplicity-corrected Šidák's tests. n=8. **D)**

Weekly average total fluid intake (mL/day) for each mouse in full sample (both males and females) after unpaired 4-OHT. There were no significant differences in the weekly average of preference between the groups by multiplicity-corrected Šidák's tests. n=8. **E)** Alcohol preference after different single doses of 4-OHT in 2BC assay in female mice. 2-way RM ANOVA omnibus effects: $F_{Group}(4,26) = 1.887$, $p=0.14$; $F_{GroupxTime}(10.24,66.57) = 1.957$, $p=0.051$; n=6-7 (all females). **F)** Weekly average alcohol preference after different single doses of 4-OHT in 2BC assay in female mice. One-way ANOVA on week 2: $F(4,26) = 4.026$, $p=0.011$; $^{**}p<0.01$, $^{*}p<0.05$ by Holm-Šidák *post-hoc* test adjusted for multiple comparisons. n=6-7 (all females). **G)** Alcohol preference weekly average for visualization of individual data points in full sample (both males and females) of mice in 2BC assay after tamoxifen administration. Asterisks represent differences between vehicle group and 2mg/kg tamoxifen group (light red) and 8mg/kg tamoxifen group (dark red) by multiplicity-corrected Šidák tests after 2-way RM ANOVA. $^{****}p<0.0001$, $^{***}p<0.001$, $^{**}p<0.01$, $^{*}p<0.05$. n=7-8. **H)** Alcohol intake weekly average (g/kg/24h) for visualization of individual data points in full sample (both males and females) of mice in 2BC assay after tamoxifen administration. Asterisks represent differences between vehicle group and 2mg/kg tamoxifen group (light red) and 8mg/kg tamoxifen group (dark red) by multiplicity-corrected Šidák tests after 2-way RM ANOVA. $^{***}p<0.001$, $^{**}p<0.01$. n=7-8. **I)** Total fluid intake (mL/day) in full sample (both males and females) of mice in 2BC assay after tamoxifen administration. 2-way RM ANOVA omnibus effects: $F_{Group}(2,20) = 1.442$, $p=0.26$; $F_{GroupxTime}(14.83,148.3) = 1.168$, $p=0.30$. No significant effects in pre-treatment period. n=7-8. **J)** Total fluid intake weekly average (mL/day) for visualization of individual data points in full sample (both males and females) of mice in 2BC assay after tamoxifen administration. There were no significant differences in the weekly average of preference between the groups by multiplicity-corrected Šidák's tests. n=7-8. All data shown as mean ± SEM.

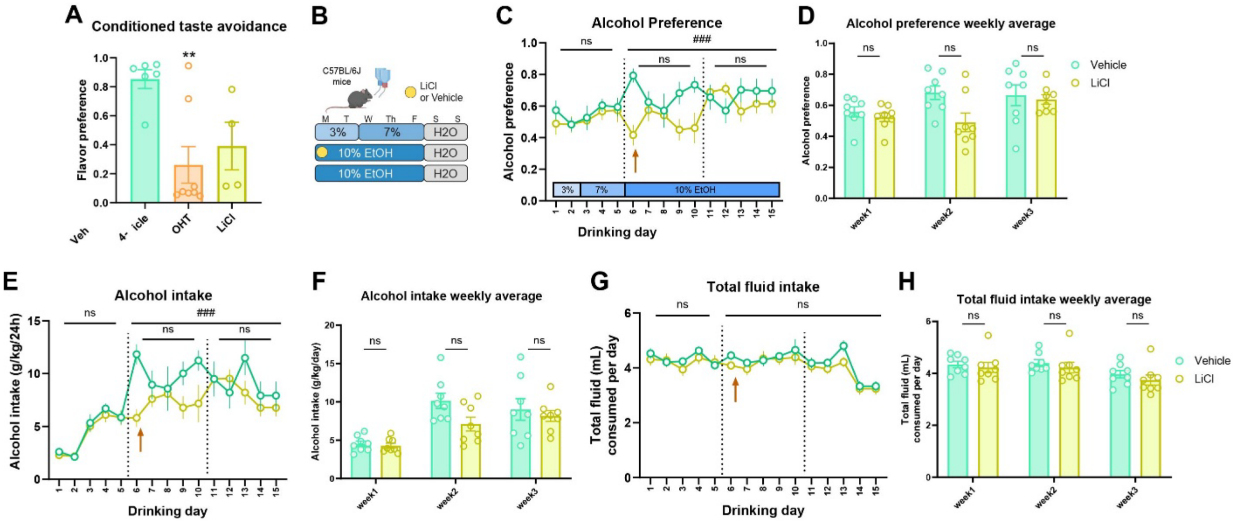


**Extended Data Figure 5. Conditioned taste avoidance (CTA) behavior and alcohol preference and consumption after single dose of lithium chloride (LiCl)**. **A)** Flavor preference following CTA for a single dose of 4-hydroxytamoxifen (4-OHT; 50mg/kg) and LiCl (125mg/kg). Kruskal-Wallis test $\chi^2(2) = 8.37$, $p$=0.008; ** $p$<0.01 by Dunn's *post-hoc* test; n=4-8. **B)** Experimental schematic of C57BL/6J mice (n=8; 4 each males/females in LiCl group and 5 males/3 females in vehicle group) given a single dose of 125mg/kg of LiCl i.p. on the first day of 10% alcohol consumption. prior to the introduction of escalating alcohol concentrations. **C)** Alcohol preference in full sample (both males and females) of mice in 2BC assay after LiCl administration. Arrow indicates day of LiCl treatment. 2-way RM ANOVA omnibus effects in post-treatment period: $F_{Group}(1,14) = 2.497$, $p$=0.14; ###$F_{GroupxTime}(9,126) = 4.091$, $p$=0.0001; n=8. There were no significant differences in the weekly average of preference between the groups by multiplicity-corrected Šidák's tests. No significant effects in pre-treatment period. **D)** Alcohol preference weekly average for visualization of individual data points in full sample (both males and females) of mice in 2BC assay after LiCl administration. As noted in C), there were no significant differences in the weekly average of preference between the groups by multiplicity-corrected Šidák's tests. **E)** Alcohol intake (g/kg/24h) in full sample (both males and females) of mice in 2BC assay after LiCl administration. $F_{Group}(1,14) = 1.933$, $p$=0.19; ###$F_{GroupxTime}(9,126) = 3.523$, $p$=0.0006; n=8. There were no significant differences in the weekly average intake between the groups by multiplicity-corrected Šidák's tests. No significant effects in pre-treatment period. **F)** Weekly average alcohol intake (g/kg/24h) for visualization of individual data points in full sample (both males and females) of mice in 2BC assay after LiCl administration. As noted in E), there were no significant differences in the weekly average of intake between the groups by multiplicity-corrected Šidák's tests. **G)** Total fluid intake (mL/day) in full sample (both males and females) of mice in 2BC assay after LiCl administration. $F_{Group}(1,14) = 0.7127$, $p=0.41$; $F_{GroupxTime}(9,126) = 0.8948$, $p=0.53$; n=8. No significant effects in pre-treatment period. **H)** Weekly average total fluid intake (mL/day) for each mouse in full sample (both males and females) of mice in 2BC assay after LiCl administration. As noted in G), there were no significant differences either at the omnibus level or in the weekly average of intake between the groups by multiplicity-corrected Šidák's tests. All data shown as mean ± SEM.

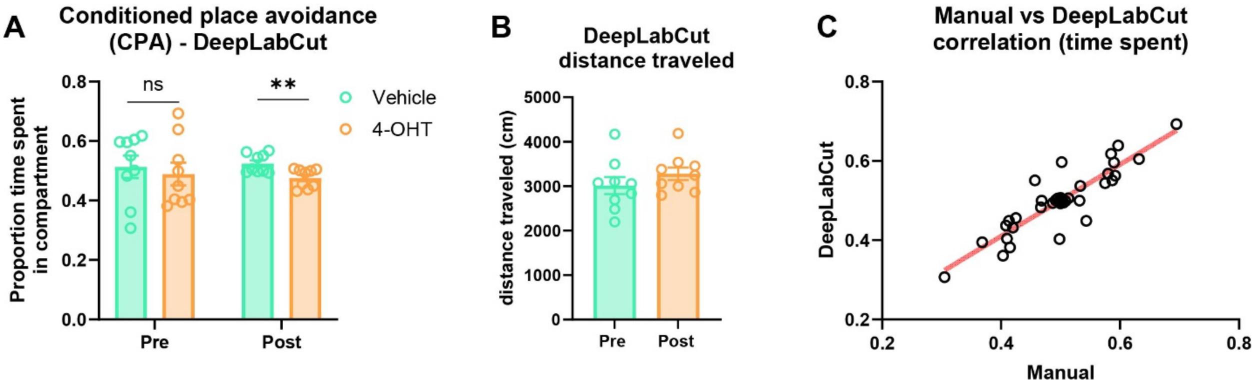


**Extended Data Figure 6. Semi-automated analysis of conditioned place avoidance (CPA)**. **A)** CPA assay before (pre) and after (post) 3 conditioning sessions each with vehicle or 50mg/kg 4-hydroxytamoxifen (4-OHT; i.p.) analyzed using DeepLabCut. $p$<0.01 by unpaired t-test; n=9. Data shown as mean ± SEM. **B)** Distance traveled analyzed in DeepLabCut. No significant effects by paired t-test; n=9. Data shown as mean ± SEM. **C)** Correlation in proportion of time spent in each compartment between manual and semi-automated (DeepLabCut) analyses. Pearson's $r$=0.8856; $p$<0.0001; n=36 pairs of observations.

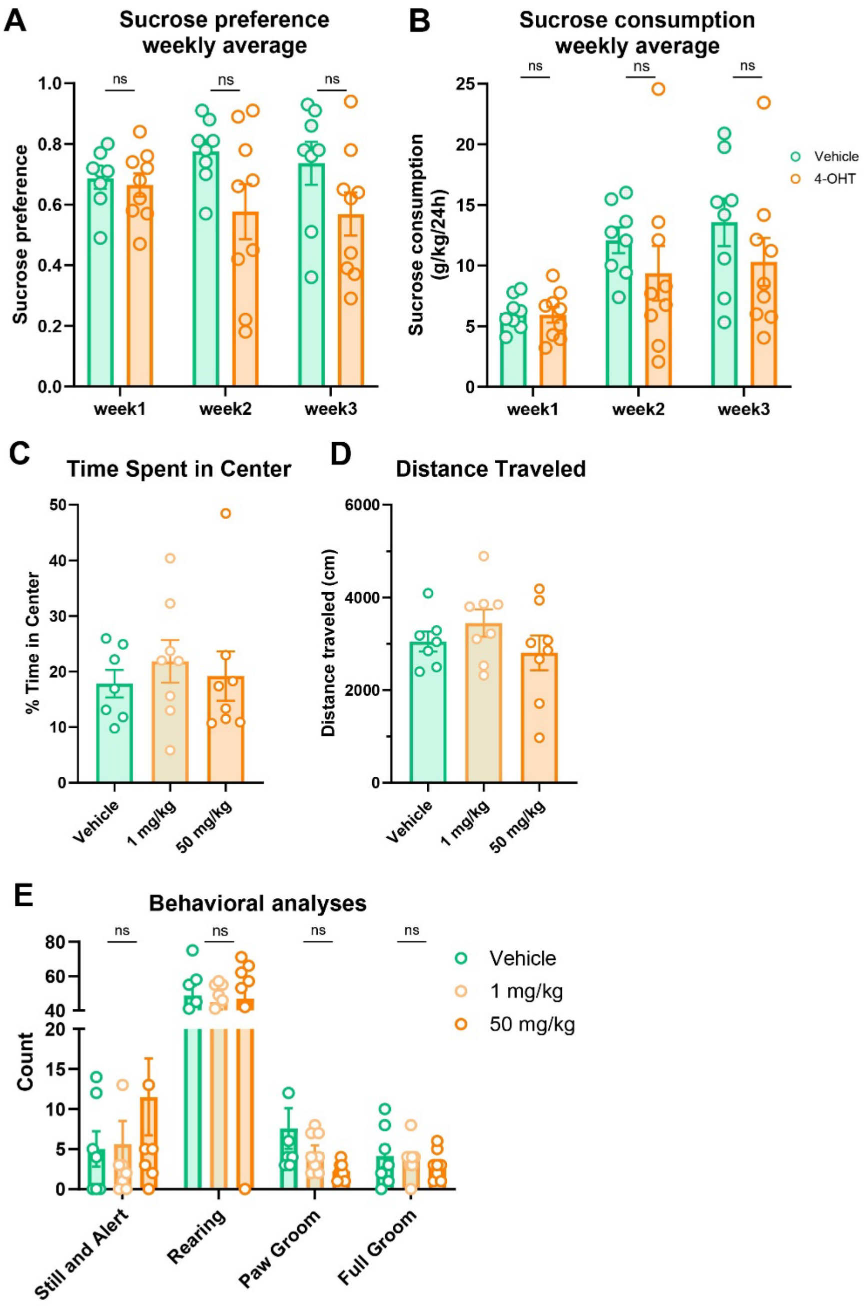


**Extended Data Figure 7. Sucrose consumption and open field behavior after 4-hydroxytamoxifen (4-OHT)**. **A)** Sucrose preference weekly average for visualization of individual data points in full sample (both males and females) of mice given 4-OHT (50mg/kg) on the first day of 1% sucrose exposure in sucrose preference test (SPT). As reported in the main figure, there were no significant differences between the groups by multiplicity-corrected Šidák's tests; n=8-9. **B)** Sucrose intake weekly average (g/kg/24h) for visualization of individual data points in full sample (both males and females) of mice given 4-OHT (50mg/kg) on the first day of 1% sucrose exposure in SPT. As reported in the main figure, there were no significant differences between the groups by multiplicity-corrected Šidák's tests; n=8-9. **C)** Time spent in center zone of open field test (OFT) in Trap2:Ai14 mice given either 1mg/kg or 50mg/kg of 4-OHT. Not significant by one-way ANOVA: $F(2,20) = 0.2880$, $p=0.75$, n=7-8. **D)** Distance traveled (cm) in OFT in Trap2:Ai14 mice given either 1mg/kg or 50mg/kg of 4-OHT. Not significant by one-way ANOVA: $F(2,20) = 1.142$, $p=0.34$, n=7-8. **E)** Behavioral phenotyping in OFT in Trap2:Ai14 mice given either 1mg/kg or 50mg/kg of 4-OHT. 2-way ANOVA omnibus effects: $F_{Group}(2,80) = 0.1932$, $p=0.82$; $F_{GroupxTime}(6,80) = 0.4799$, $p=0.82$. No significant group-level effects by multiplicity-corrected Šidák's tests. ns = not significant. All data shown as mean ± SEM.

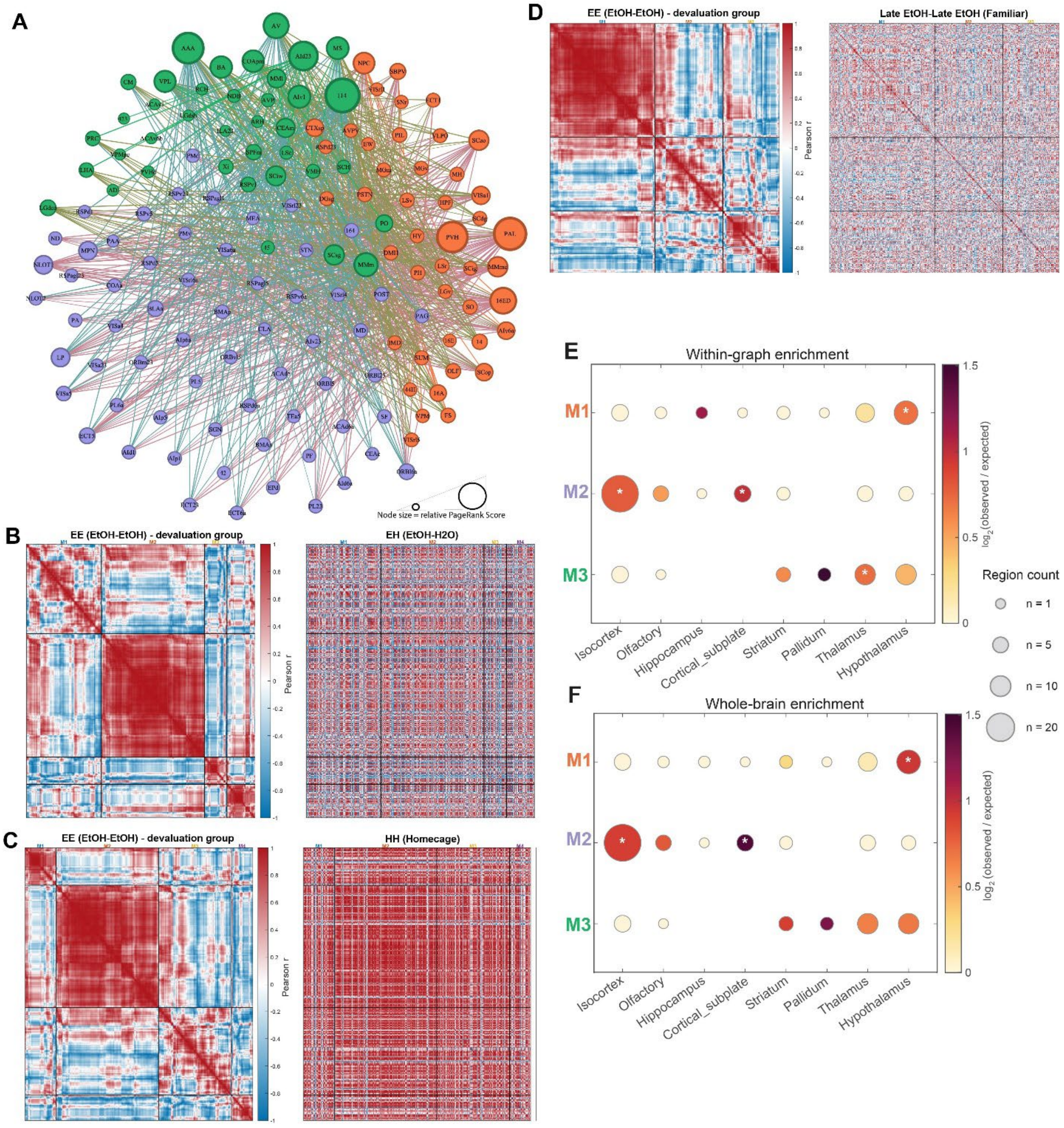


**Extended Data Figure 8. Additional graph theoretical measures for an activity screen across the brain during 4-OHT-mediated alcohol devaluation**. **A)** Negatively-correlated network connectivity in the alcohol devaluation (EE) group (n=5). Each line indicates a negative correlation (R<-0.6) between brain areas [Fruchterman-Reingold layout]; Colored nodes indicate module assignment, and the size of the node indicates the PageRank score. **B)** Comparison of correlation matrices between EE devaluation mice and mice exposed to alcohol at timepoint 1 (i.e., in conjunction with 4-OHT) but water at timepoint 2 (EtOH-H2O or EH group). Hierarchical clustering in the EE group determined the location of brain areas on both matrices. Mantel test for matrix dissimilarity (Spearman $r$): 0.0496; $p=0.006$. **C)** Same as C) but comparing EE group and a group of mice exposed to both timepoints in the homecage (HH group). Mantel test: 0.1576; $p<0.0002$. **D)** Same as C) but comparing EE group and a group of mice exposed to 4-OHT in conjunction with 10% alcohol after 6 weeks of drinking experience (for timepoint 1) and then sacrificed while drinking alcohol 1 week later (late EtOH-late EtOH or LL group). Mantel test: 0.0511; $p<0.0002$. Note that small differences in the brain region clustering order for the EE matrix result from clustering iterations. **E)** Dot plot showing enrichment for module membership based on aggregate brain areas in Allen Brain Atlas using only areas input into the graph (significant areas from TTP-S) as the denominator for enrichment. Dot size indicates the number of areas in the module, and dot color indicates enrichment as log2(observed/expected). *$p<0.05$ by Fisher's exact test. **F)** Same as E) using all areas tested for significance as the denominator. *$p<0.05$ by Fisher's exact test.

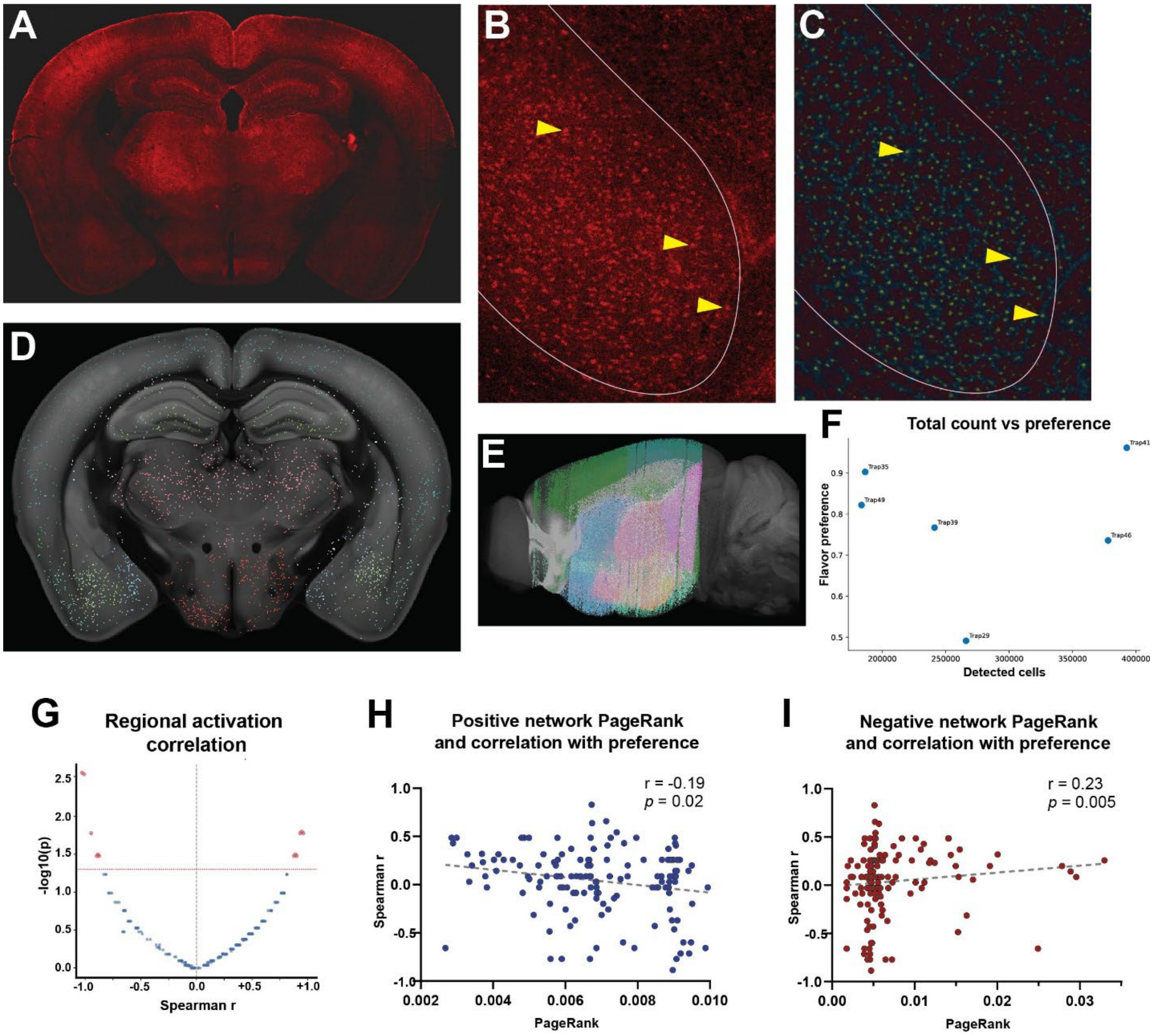


**Extended Data Figure 9. Expression and quantification of mCherry (AAV-PHP.eb-hSyn-DIO-hM3D(Gq)-mCherry) in brain following peripheral injection.** Trap2 mice were given a retroorbital injection of a CNS-penetrant chemogenetic construct (AAV-PHP.eb-hSyn-DIO-hM3D(Gq)-mCherry) prior to activation of a simple homecage 'Trap' ensemble via injection of 4-OHT (50mg/kg; i.p.) 3 weeks later. After 4 weeks to allow for recombination, animals completed a CTA, and brains were harvested and perfused for immunohistochemistry. **A)** Example brain slice to visualize mCherry expression at 4x. **B)** Zoom of basolateral amygdala showing staining of cells, projections, and background. Neurons are denoted with yellow arrows. **C)** Ilastik training images showing transfected neurons in yellow, projections in dark blue, and background in dark red/black. The same neurons from B) are denoted with yellow arrows. **D)**

Quantification image from BrainJ showing detected cells in an example brain slice mapped to the Allen Brain Atlas Common Coordinate Framework (CCFv3). **E)** Three-dimensional rendering of detected mCherry-expressing neurons in an example brain. **F)** Correlation between total detected mCherry-positive neurons and flavor preference. (Spearman $r$=-0.03; $p$=0.957). **G)** Volcano plot of Spearman r correlation coefficient of expression of mCherry/hM3D(Gq) from each of ~500 brain areas with flavor preference shown. The 12 regions highlighted in red demonstrated uncorrected p-values below 0.05, and no single region met multiple comparison correction criteria for significance. Cell counts were corrected for regional volume. **H)** Correlation between PageRank from positive correlation network in alcohol devaluation group and correlation between flavor preference in homecage Trap CTA and brain region-specific mCherry expression. Regions with a higher PageRank score in the alcohol devaluation network tend be correlated with flavor avoidance in a separate experiment (Spearman $r$= -0.19; $p$=0.02). **I)** Same as H) but for PageRank with negative correlation network. Regions with a higher PageRank score in the alcohol devaluation network tend be anticorrelated with flavor avoidance in a separate experiment (Spearman $r$= 0.23; $p$=0.005).

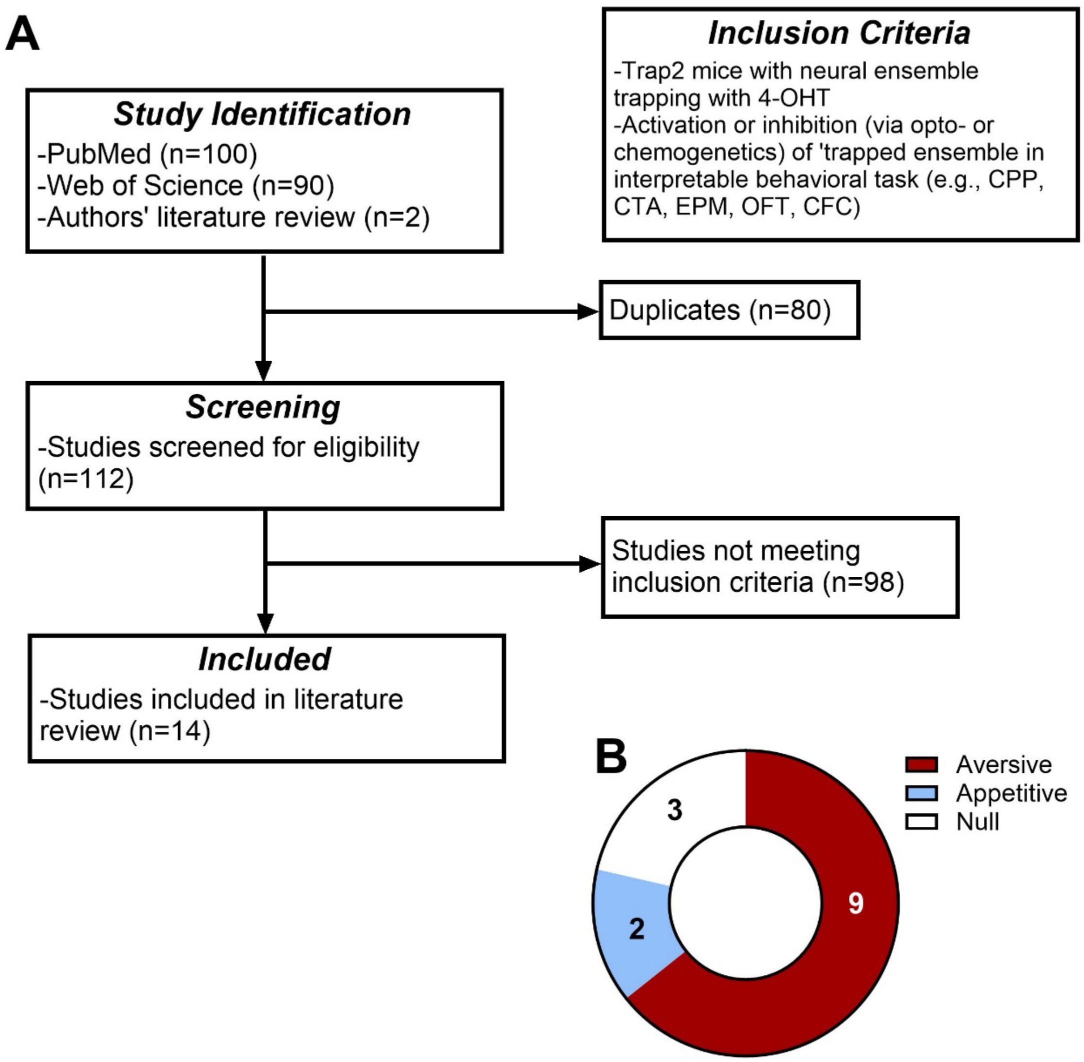


**Extended Data Figure 10. Systematic review of effects of re-activating 'trapped' ensembles**. **A)** Diagram of review design. See Methods for search terms and further details on inclusion and exclusion criteria. **B)** Included studies were categorized based on whether re-activation of the 'trapped' ensemble led to aversive effects (e.g., avoidance in conditioned taste/place avoidance assays; thigmotaxis in open field test), appetitive effects (e.g., increased time spent in paired compartment in conditioned place preference; increased exploration of open arms in elevated plus maze), or no significant effects (null).
